# Schizophrenia Exhibits Bi-Directional Brain-Wide Alterations in Cortico-Striato-Cerebellar Circuits

**DOI:** 10.1101/166728

**Authors:** Jie Lisa Ji, Caroline Diehl, Charles Schleifer, Carol A. Tamminga, Matcheri S. Keshavan, John A. Sweeney, Brett A. Clementz, S. Kristian Hill, Godfrey Pearlson, Genevieve Yang, Gina Creatura, John H. Krystal, Grega Repovs, John Murray, Anderson Winkler, Alan Anticevic

**Author notes:** **Corresponding Author:** Alan Anticevic, Yale University, Department of Psychiatry 40 Temple St, Suite 6E, New Haven, CT 06520.

## Abstract

Distributed neural dysconnectivity is considered a hallmark feature of schizophrenia, yet a tension exists between studies pinpointing focal disruptions versus those implicating brain-wide disturbances. The cerebellum and the striatum communicate reciprocally with the thalamus and cortex through monosynaptic and polysynaptic connections, forming cortico-striatal-thalamic-cerebellar (CSTC) functional pathways that may be sensitive to brain-wide dysconnectivity in schizophrenia. It remains unknown if the same pattern of alterations persists across CSTC systems, or if specific alterations exist along key functional elements of these networks. We characterized connectivity along major functional CSTC subdivisions using resting-state functional magnetic resonance imaging in 159 chronic patients and 162 matched controls. Associative CSTC subdivisions revealed consistent brain-wide bi-directional alterations in patients, marked by hyper-connectivity with sensory-motor cortices and hypo-connectivity with association cortex. Focusing on the cerebellar and striatal components, we validate the effects using data-driven *k*-means clustering of voxel-wise dysconnectivity and support vector machine classifiers. We replicate these results in an independent sample of 202 controls and 145 patients, additionally demonstrating that these neural effects relate to cognitive performance across subjects. Taken together, these results from complementary approaches implicate a consistent motif of brain-wide alterations in CSTC systems in schizophrenia, calling into question accounts of exclusively focal functional disturbances.

## INTRODUCTION

A fundamental challenge in clinical neuroscience is the search for robust neuroimaging biomarkers reflecting disease-related alterations in large-scale neural systems. Advances in human neuroscience offer opportunities for clinical translations and biomarker development. For instance, emerging human neuroimaging studies have identified the presence of large-scale cortico-striatal-thalamic-cerebellar (CSTC) functional pathways that are stable and replicable across hundreds of individuals (Buckner RL et al. 2011; Yeo BT et al. 2011; Choi EY et al. 2012). Such pathways may be particularly sensitive to disruptions in psychiatric illnesses such as schizophrenia (SCZ), which has been conceptualized as a disorder of synaptic communication affecting distributed neural systems. Leading theoretical models of SCZ propose that disruptions in local microcircuit excitation (E) and inhibition (I) neuronal balance, due to altered synaptic signaling, may underlie noted alterations across neural systems (Stephan KE et al. 2006; Murray JD, A Anticevic, et al. 2014; Yang GJ, JD Murray, XJ Wang, et al. 2016). One possibility is that such alterations occur only in select CSTC functional pathways if such pathways are more vulnerable to the upstream microcircuit perturbations. Alternatively, such synaptic perturbations may generalize across distributed CSTC pathways. Hence, a major gap in knowledge concerns whether neural disturbances in SCZ manifest brain-wide across CSTC functional pathways, or if they occur in only select regions of these circuits. Addressing this tension is critical to informing neuro-marker development that can guide treatment for either targeted neuropathology in specific areas or diffuse alterations that span CSTC pathways.

One widely replicated effect in SCZ neuroimaging is the disruption in functional relationships involving the thalamic resting-state blood-oxygen-level-dependent (BOLD) signal (Ramsay IS and AW MacDonald, 3rd 2018) The thalamus consists of topographically organized nuclei that are densely connected with other brain areas (Ray JP and JL Price 1993; Haber S and NR McFarland 2001; Klein JC et al. 2010), making it highly sensitive to widespread disturbances neural disturbances. Specifically, several independent replications have established a bi-directional pattern of dysconnectivity with the thalamus, characterized by elevated thalamic coupling with sensorimotor brain regions and reduced thalamic coupling with associative cortical areas (Welsh RC et al. 2010; Woodward ND et al. 2012; Anticevic A, MW Cole, et al. 2014). These effects generalize across patients with chronic SCZ and prodromal individuals at ultra-high-risk for developing SCZ (Anticevic A et al. 2015). Critically, these effects are most pronounced along higher-order associative thalamic functional subdivisions (e.g. mediodorsal nucleus). Multiple theoretical models have proposed that these disruptions in thalamic functional coupling may be a hallmark of the illness (Lisman JE et al. 2010). Yet, it remains unknown if these disrupted patterns manifest across the entire CSTC functional pathway or if they are exclusive to the thalamus. If the thalamic effect is indeed specific, then similar patterns would not be expected to emerge across other CSTC systems as robustly. However, if SCZ involves synaptic neural alterations that are not exclusive to the thalamus and affect other CSTC components, then we hypothesize that the described bi-directional effects may emerge across brain-wide CSTC systems, reflecting shared disruption. Previous studies have identified alterations in isolated regions of the basal ganglia and cerebellum in SCZ (Andreasen NC et al. 1996; Holt DJ et al. 2005; Konarski JZ et al. 2005; Rusch N et al. 2007; Collin G et al. 2011; Liu H et al. 2011; Tu PC et al. 2012; Wang L et al. 2014; Sarpal DK et al. 2015; Sarpal DK et al. 2016), but to date none have examined whether disturbances in functional relationships are unified across these neural systems.

A way to close this knowledge gap involves leveraging existing resting-state findings in healthy humans that have defined large-scale functional networks across cortical, striatal and cerebellar neural territories (Buckner RL *et al.* 2011; Yeo BT *et al.* 2011; Choi EY *et al.* 2012). This work implicated a shared functional architecture across the CSTC systems that can be used to assay patterns of changes across each functional CSTC subdivision in SCZ. We focus here on the striatum and cerebellum, as using the striatal or cerebellar elements of each functional network allows us to characterize altered functional relationships across CSTC systems while bypassing the thalamus as a starting ‘seed’ point. Notably, the cerebellum does not share monosynaptic anatomical connections with cortex, thalamus or striatum (hence is not a part of the anatomically-defined cortico-striatal-thalamic-cortical loop). However, it is highly functionally connected these areas via multiple pathways, as evidenced by stable and cohesive large-scale functional networks across these neural territories (Buckner RL *et al.* 2011; Yeo BT *et al.* 2011; Choi EY *et al.* 2012). Hence the cerebellum serves as an important ‘control’ in our study: if cerebellar effects are seen, it would support the hypothesis that disruptions in SCZ are pervasive across functional brain-wide networks, and not limited by thalamic or striatal anatomy *per se*. Even if such patterns are revealed, it is unknown known if these ‘bidirectional’ dysconnectivity CSTC effects may relate to psychosis symptoms or cognitive deficits. Prior work has demonstrated that thalamic dysconnectivity may present a robust cognitive remediation target in schizophrenia (Ramsay IS et al. 2017). Therefore, we tested the hypothesis that, if observed, CSTC dysconnectivity may be predictive of cognitive impairment in patients.

In summary, we report robust bi-directional CSTC dysconnectivity patterns that are prominent across the cerebellar and striatal functional subdivisions, including the thalamus and cortex. This CSTC dysconnectivity in SCZ is inconsistent with the possibility of exclusive thalamic disruptions. Results replicated and revealed most pronounced alterations along higher-order associative divisions of the CSTC networks, using both an independently defined *a priori* parcellation and a data-driven clustering approaches. The severity of CSTC dysconnectivity was related to cognition but not psychosis symptoms. Collectively, these effects implicate distributed associative CSTC alterations in SCZ in support of brain-wide disruptions in information flow that may preferentially relate to severity of cognitive deficits.

## MATERIALS AND METHODS

### Primary Dataset

For the primary analyses, we analyzed data from 159 patients with chronic schizophrenia and 162 demographically matched healthy controls (**Table 1**) recruited from the Olin Neuropsychiatry Research Center and from a publicly distributed dataset provided by the Center for Biomedical Research Excellence (COBRE). All subjects met identical methodological stringency criteria and underwent identical analyses. Of note, because each independent sample included both patients and matched controls, we collapsed the two samples for our analyses. Additional details are given in **Supplementary Materials and Methods**.

**Table 1.**
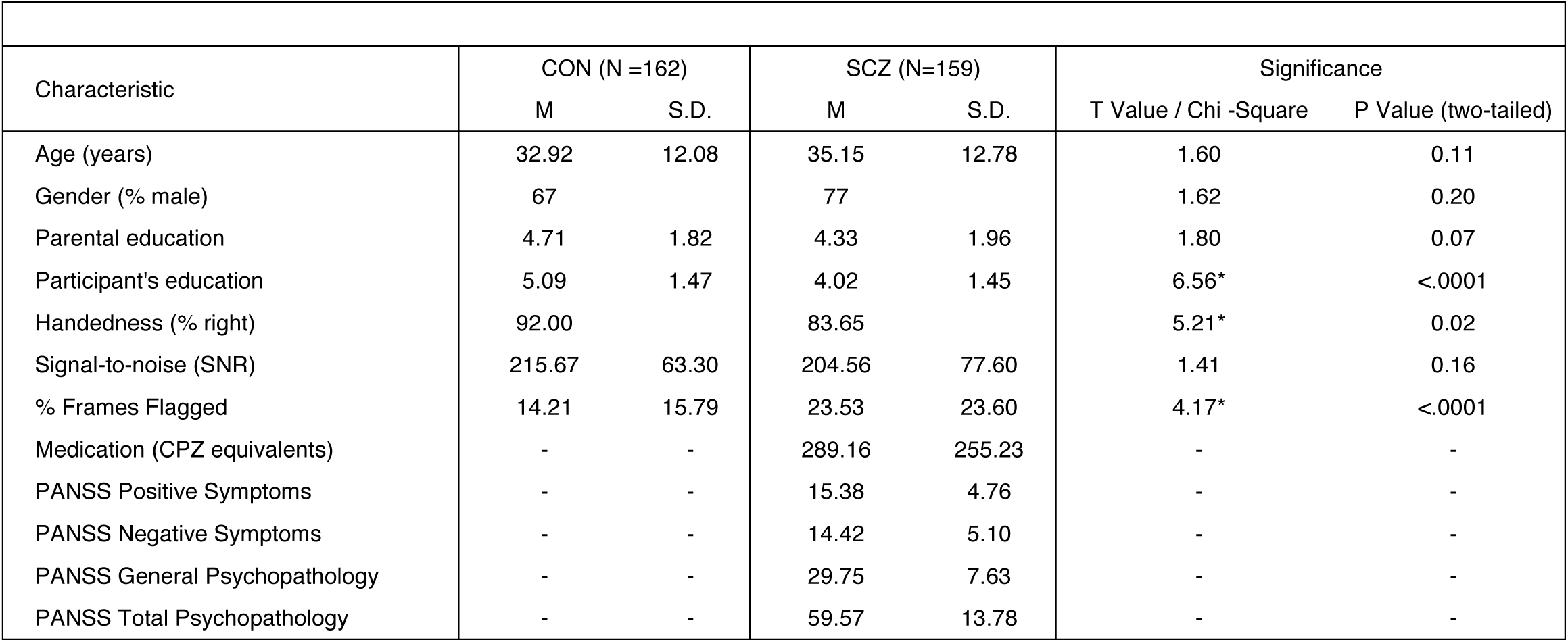
Demographics for discovery sample. CON, control subjects; SCZ, Patients diagnosed with schizophrenia; PANSS, Positive and Negative Syndrome Scale; M, Mean; SD, Standard Deviation. Education level for the COBRE sample was determined based on the following scale: Grade 6 or less=1; Grade 7–11=2; high school graduate=3; attended college=4; graduated 2-year college=5; graduated 4-year college=6; attended graduate or professional school=7; Completed graduate or professional school=8. Olin education data were converted from ‘years of education’ to the COBRE education scale in order to permit combining of education demographic data. Parental education for the Olin set is the average of the mother and father’s education. CPZ, Chlorpromazine equivalents were calculated according to latest validated approaches (Andreasen NC *et al.* 2010). SNR (signal-to-noise ratio) was determined by obtaining the mean signal and standard deviation for a given slice across the relevant BOLD run, while excluding all non-brain voxels across all frames (Anticevic A, G Repovs*, et al.* 2012). An ^*^ denotes a significant T statistic for the between-group t-test, uncorrected for multiple comparisons.

| Table 1 \| Discovery Sample - Clinical and Demographic Characteristics |  |  |  |  |  |  |
| --- | --- | --- | --- | --- | --- | --- |
| Characteristic | CON (N =162) |  | SCZ (N=159) |  | Significance |  |
|  | M | S.D. | M | S.D. | T Value / Chi -Square | P Value (two-tailed) |
| Age (years) | 32.92 | 12.08 | 35.15 | 12.78 | 1.60 | 0.11 |
| Gender (% male) | 67 |  | 77 |  | 1.62 | 0.20 |
| Parental education | 4.71 | 1.82 | 4.33 | 1.96 | 1.80 | 0.07 |
| Participant's education | 5.09 | 1.47 | 4.02 | 1.45 | 6.56* | <.0001 |
| Handedness (% right) | 92.00 |  | 83.65 |  | 5.21* | 0.02 |
| Signal-to-noise (SNR) | 215.67 | 63.30 | 204.56 | 77.60 | 1.41 | 0.16 |
| % Frames Flagged | 14.21 | 15.79 | 23.53 | 23.60 | 4.17* | <.0001 |
| Medication (CPZ equivalents) | - | - | 289.16 | 255.23 | - | - |
| PANSS Positive Symptoms | - | - | 15.38 | 4.76 | - | - |
| PANSS Negative Symptoms | - | - | 14.42 | 5.10 | - | - |
| PANSS General Psychopathology | - | - | 29.75 | 7.63 | - | - |
| PANSS Total Psychopathology | - | - | 59.57 | 13.78 | - | - |

### Replication Dataset

The replication and extension sample was obtained from the publicly available Bipolar-Schizophrenia Network on Intermediate Phenotypes (B-SNIP) dataset in collaboration with the BSNIP Principal Investigators. B-SNIP participants were collected across 6 different sites in the United States. Full details on the B-SNIP data acquisition and characterization of site effects have been previously rigorously characterized (Tamminga CA et al. 2013; Meda SA et al. 2015; Sheffield JM et al. 2017). Briefly, subjects were excluded if they had i) a history of seizures or head injury resulting in >10 minutes loss of consciousness, ii) positive drug screen on the day of testing, iii) a diagnosis of substance abuse in the past 30 days or substance dependence in the past 6 months, iv) history of serious medical or neurological disorder that would likely affect cognitive functioning, v), history of serious medical or neurological disorder that would likely affect cognitive functioning, vi) insufficient English proficiency, or vii) an age-corrected Wide-Range Achievement Test (4th edition) reading test standard score <65. Additionally, patients were required to have had no change in medication and been clinically stable over the past month. Demographic information for this dataset is given in **Supplementary Table S1**. Details on acquisition parameters across the 6 sites are given in **Supplementary Table S2**. After preprocessing and quality control (described below), data from 145 patients with schizophrenia and 202 healthy controls were included. Critically, participants in the B-SNIP dataset underwent the Brief Assessment of Cognition in Schizophrenia (BACS) battery, which provided an assessment of cognitive functioning (Keefe RS et al. 2004). The composite scores used here are presented as standardized Z-scores normalized to mean and standard deviation of the control group and were used to test the hypothesis that, if observed, CSTC dysconnectivity may be predictive of cognitive impairment in patients.

### Symptoms and Medication

The severity of schizophrenia symptoms was assessed using the Positive and Negative Syndrome Scale (PANSS) (Kay SR et al. 1987), which includes positive, negative, and general psychopathology symptom dimensions (Kay SR *et al.* 1987) (**Table 1**). 75 of the 90 schizophrenia patients in the Olin sample were receiving antipsychotic treatment, and all patients in the COBRE sample were receiving stable doses of antipsychotic medication with no medication changes for at least 1 month. All medications were converted to chlorpromazine equivalents and used as covariates (Andreasen NC et al. 2010).

### Neuroimaging Data Acquisition

Neuroimaging data acquired at the Olin Neuropsychiatry Research Center were obtained using a Siemens-Allegra 3-T scanner, with axial slices parallel to the anterior–posterior commissure (AC–PC) using a T2^*^-weighted gradient-echo, echo-planar sequence [time repetition (TR)/time echo (TE)=1500/27ms, flip angle=60°, field of view=24 × 24 cm, acquisition matrix=64 × 64, voxel size=3.43 × 3.43 × 4 mm], ensuring whole-brain coverage. The acquisition lasted 5.25 min and produced 210 volumetric images per subject (29 slices/volume, inter-slice gap=1 mm). Subjects were instructed to lay awake in the scanner and keep their eyes open. Subjects were monitored on a video camera to ensure that they stayed awake and were removed from analyses if they fell asleep during the scan, or if their head movement >1 mm along any axis. Structural images were acquired using a T1-weighted, 3-dimensional magnetization-prepared rapid gradient-echo sequence (TR/TE/time to inversion=2200/4.13/766 ms, flip angle=13°, voxel size (isotropic)=0.8mm, image size=240×320×208 voxels), with axial slices parallel to the AC–PC line. Subjects that comprised the COBRE sample underwent data collection at Center for Biomedical Research Excellence using a Siemens Tim-Trio 3T scanner. Full acquisition details for the COBRE SCZ sample and CON have been detailed previously (Yang GJ, JD Murray, X-J Wang, et al. 2016). Briefly, BOLD signal was collected with 32 axial slices parallel to the AC-PC using a T2^*^-weighted gradient-echo, echo-planar sequence (TR/TE=2000/29ms, flip angle=75°, acquisition matrix=64×64, voxel size=3×3×4mm). The acquisition lasted 5 minutes and produced 150 volumetric images per subject. Structural images were acquired using a 6-minute T1-weighted, 3D MPRAGE sequence (TR/TE/TI=2530/[1.64, 3.5, 5.36, 7.22, 9.08]/900, flip angle=7°, voxel size (isotropic)=1mm, image size=256×256×176 voxels) with axial slices parallel to the AC-PC line. Details on acquisition parameters across the 6 sites comprising the replication B-SNIP sample are given in **Supplementary Table S2**.

### Data Preprocessing and Analysis

For the primary dataset, all preprocessing followed previously validated approaches (Repovs G et al. 2011; Anticevic A, M Gancsos, et al. 2012; Anticevic A, G Repovs, et al. 2012; Repovs G and DM Barch 2012) to ensure continuity with reported thalamic effects. Critically, the B-SNIP ‘replication’ dataset was preprocessed in accordance with the Human Connectome Project (HCP) minimal preprocessing pipeline (Glasser MF et al. 2013). We made the choice to use HCP pipelines for the replication dataset (as opposed to processing methods in the primary dataset following published procedures) to ensure generalizability of effects across pipelines choices. This allowed us to verify two key observations: i) if the prior thalamic generalizes across the CSTC systems using prior methods; ii) if the effects replicate and remain robust to preprocessing pipeline choices. Details on the HCP adaptation for ‘legacy’ B-SNIP data are comprehensively described in the **Supplement.**

Irrespective of pipeline choice, all data underwent consistent standard practices processing procedures. BOLD data underwent: i) slice-time correction, ii) first 5 images removed from each run, iii) rigid body motion correction, iv) transform of the structural images to the standard template, and v) co-registration of BOLD volumes to the standard structural image template. All structural data underwent FreeSurfer’s recon-all pipeline to compute brain-wide segmentation of gray and white matter, which was used to define anatomical nuisance regressors and brain-wide grey matter masks. Additionally, all BOLD images had to pass stringent quality assurance criteria to ensure that all functional data were of comparable and high quality. We excluded images with signal-to-noise ratios (SNR)<90, computed by obtaining the mean signal and standard deviation (SD) for a given slice across the BOLD run, while excluding all non-brain voxels across all frames (Anticevic A, MS Brumbaugh, et al. 2012) (see **Table 1**). To remove sources of spurious correlations present in resting-state BOLD data, all time-series were high-pass filtered (>0.08Hz). The following regressors were used in additional BOLD de-noising: i) nuisance signal removal from ventricles, ii) deep white matter, iii) global grey matter mean signal (GMS), iv) 6 rigid-body motion correction parameters, and their first derivatives. Finally, data were additionally low-pass temporal filtered (<0.009Hz). In addition, we implemented “movement scrubbing” as recommended by (Power JD et al. 2012). Movement scrubbing refers to the practice of removing BOLD volumes that have been flagged for high motion in order to minimize movement artifacts. Specifically, all frames with possible movement-induced artifactual fluctuations in intensity were identified via two criteria: i) frames in which the sum of the displacement across all 6 rigid body movement correction parameters exceeded. 5mm (assuming 50mm cortical sphere radius) were identified; ii) root mean square (RMS) of differences in intensity between the current and preceding frame was computed across all voxels divided by mean intensity and normalized to time series median. Frames in which normalized RMS exceeded the value of 3 were identified. The frames flagged by either criterion were marked for exclusion (logical or), as well as the one preceding and two frames following the flagged frame. Importantly, levels of motion and SNR did not relate to reported effects (**Supplementary Fig. S7-S9**).

### Network and Parcel Terminology

Throughout our analyses we make a distinction between the sensory networks (VIS, visual; SOM, somatomotor) and the associative networks (DAN, dorsal attention; VAN, ventral attention; LIM, limbic; FPCN, frontoparietal control; and DMN, default mode). We use the term “parcel” to refer to sets of voxels in the cerebellum or striatum that are defined by their membership to one of the functional networks. For instance, the cerebellar FPCN parcel is the set of voxels that comprises the FPCN network subdivision in the cerebellum. Put differently, the FPCN network refers to the brain-wide set of voxels that belong to this network, whereas the cerebellum FPCN “parcel” is the set of voxels within the cerebellum that belong to the FPCN, as defined by Buckner and colleagues. Similarly, the striatum FPCN “parcel” is the set of striatal voxels that belong to the FPCN, as defined by Choi and colleagues.

### Seed-Based Functional Connectivity (fcMRI) Analyses

Throughout, parallel analyses were conducted independently for the cerebellum and the striatum parcels. In-house tools (Repovs G *et al.* 2011) were used to compute whole-brain correlation maps by extracting average time-series across all voxels within a given subject for each of the 7 cerebellar and 6 striatal parcels as defined by Buckner et al. (2011) and Choi et al. (2012). Importantly, we *did not* seed each possible isolated region from the cerebellar and striatal solution, but rather treated all the FPCN-belonging voxels in the cerebellum (i.e. cerebellum FPCN parcel) as a ‘seed’. The same logic applies to the striatum.

This average striatal or cerebellar signal for each element/parcel was then correlated with each gray matter voxel in the brain. The computed Pearson correlation values were transformed to Fisher Z-values (Fz) using a Fisher’s *r*-to-*Z* transform. This yielded 7 cerebellar and 6 striatal whole-brain Fz maps for each subject, one for each parcel seed. For every such map each voxel’s value represents connectivity with the original cerebellar or striatal parcel.

We then computed group average connectivity maps for each parcel for each group. To test the central hypotheses, the 7 cerebellum parcel-seeded maps were entered into a 2×7 mixed model ANOVA with one between-group factor (SCZ vs. CON) and one within-subject factor (7 cerebellar parcels). We evaluated whole-brain type I error protected effects using nonparametric techniques implemented with the Permutation Analysis of Linear Models package (PALM (Smith SM and TE Nichols 2009; Winkler AM et al. 2014)). The entire general linear model was permuted 2,000 times with tail-approximation (Winkler AM et al. 2016) to obtain null distributions for every test. To circumvent the need to define clusters using arbitrary thresholds for cluster size, we used threshold-free cluster enhancement (TFCE (Smith SM and TE Nichols 2009)). Resulting statistical images were then thresholded at whole-brain corrected *p*<0.05 (controlled for family-wise error). A similar analysis was performed for striatum parcel-seeded maps but with a 2×6 mixed model ANOVA design given that the striatum contained 6 parcels. Results were visualized using the Caret 5.5 software (http://brainvis.wustl.edu/wiki/index.php/Caret), NeuroLens software (http://www.neurolens.org) and the Connectome Workbench software (http://www.humanconnectome.org/software/connectome-workbench.html) www.humanconnectome.org/software/connectome-workbench.html).

### Quantifying the Overlap Between Cerebellar and Striatal Main Effects of Group

We performed a conjunction analysis by testing for regions that were significantly hyper-or hypo-connected (*p<*0.05) in both the cerebellar and the striatal main effects of parcel (**Fig. 2c & d**). We tested for the significance of such overlap using the hyper-geometric test, given the total number of voxels in the brain and the number of significant voxels in the striatal and cerebellar maps (referred to as Map 1 and Map 2 below, as the order is arbitrary and interchangeable). We performed the test separately for the two hyper-connectivity maps and the two hypo-connectivity maps. The hypergeometric distribution describes the probability of drawing *k* successes in *n* sequential trials without replacement, in a population of size *N* containing *K* successes. Hence, the probability of obtaining *k* overlapping significant voxels between Map 1 and Map 2 can be found using the formula:

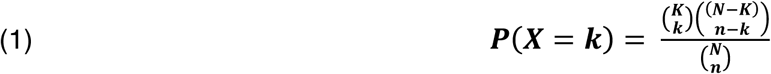

where *k* is the number of overlapping voxels, *n* is the number of significant voxels in Map 1, *K* is the number of significant voxels in Map 2, and *N* is the total number of voxels in the brain. 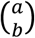 is a binomial coefficient and is calculated using:

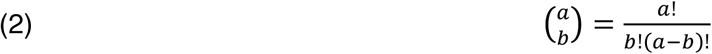

The expected value 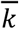 for the hypergeometric distribution, here the expected number of voxels in the overlap between Maps 1 and 2 due to chance, is:

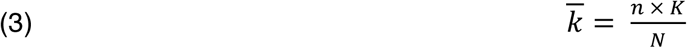

When *k* is greater than 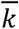, the probability of observing *k* or more voxels is obtained from the cumulative distribution function:

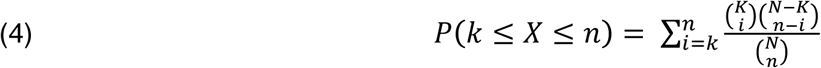

We then calculated exact 95% confidence intervals for the expected value of the hypergeometric distribution using the Clopper-Pearson approach (Clopper CJ and ES Pearson 1934). The upper limit *p*_*u*_ at confidence level *α* is given such that:

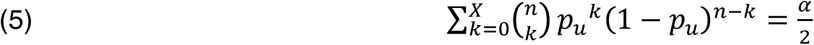

and the lower limit *p*_*l*_ is given such that:

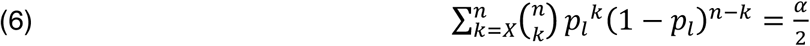

### *K*-means Clustering of Group Dysconnectivity

We first identified all grey matter voxels using FreeSurfer segmentation and computed the functional connectivity of every cerebellar voxel with all other gray matter voxels in the brain. The gray matter mask was defined based on FreeSurfer cortical segmentation codes. To ensure adequate gray matter signal, we included only voxels where at least 25% of all subjects had gray matter in the cortex and thalamus and 10% in the cerebellum and striatum (Cole MW et al. 2011; Anticevic A, MS Brumbaugh*, et al.* 2012; Anticevic A, MW Cole*, et al.* 2014). Three subjects were excluded from the clustering analyses due to incomplete coverage at the most inferior slices of the cerebellum. Next, we applied a Fisher’s *r*-to-*Z* transform to the subject-specific cerebellar/striatal connectivity maps and then computed group connectivity maps by averaging across each voxel separately for all subjects in each group. We then computed a group difference ‘dysconnectivity’ map for each cerebellar / striatal voxel, and applied *k-*means clustering on these difference maps to obtain clusters of voxels with the most similar patterns of disturbance. Specifically, 1-*r* was then used as a dissimilarity measure and the *k-*means clustering algorithm was applied to find *k* number of clusters that group together the voxels with the most similar dysconnectivity patterns across the brain. To minimize the possibility of the algorithm being caught in a local minimum, the clustering for each *k* was repeated 10 times with different random starting values and the solution with the smallest within-cluster distances was accepted (Nanetti L et al. 2009; Cauda F et al. 2011; Anticevic A, MW Cole*, et al.* 2014). See **Supplementary Fig. S22** for a schematic of the *k*-means clustering workflow. Notably, because there is no known *a priori* number of cerebellar or striatal subdivisions with distinct patterns of schizophrenia-related dysconnectivity, we examined a range of *k* values and highlight the *k=*7 solution (**Supplementary Fig. S23c** and **S26**) so that comparisons can be made with the Yeo/Buckner parcellation, which contains 7 functional networks. Alternative solutions (*k*=4 and *k=*6) are provided in **Supplementary Fig. S23a-b, S24** and **S25**. For similar reasons we highlight the *k=*6 solution for the striatum (**Supplementary Fig. S23e** and **S28**). Alternative striatal clustering solutions (*k*=4 and *k=*7) are presented in **Supplementary Fig. S23d & f, S27** and **S29**. The whole-brain connectivity maps of the *k*=7 cerebellar solution and *k*=6 striatal solution in CON subjects are provided in **Supplementary Fig. S30-S31**, illustrating the functional connectivity profiles of the obtained clusters.

### Dissimilarity in Cerebellar and Striatal Voxelwise Connectivity

We also examined the voxels in the cerebellum and striatum with the greatest difference between groups in coherence of BOLD fluctuations with the rest of the brain. Here we used the eta^2^ index, which quantifies the pattern of similarity between two signals. Eta^2^ is calculated by:

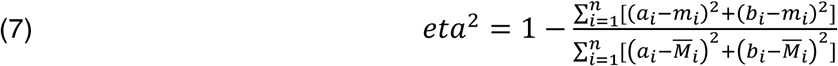

where *a*_*i*_ and *b*_*i*_ correspond to connectivity at position *i* (in this case, a given voxel in the cerebellum or striatum) for maps *a* and *b* respectively (in this case, SCZ and CON connectivity maps); *m*_*i*_ corresponds to the mean value of the two images at position *i*, i.e. (*a*_*i*_ + *b*_*i*_)/2; and 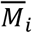 represents the grand mean value across the mean image (designated by *m*). We converted the index to 1-eta^2^ in order to show the degree of *dissimilarity* in functional connectivity in each voxel. Hence, a 1-eta^2^ value of 1 indicates no similarity between the two signals, and a 1-eta^2^ of 0 indicates perfect signal similarity. While 1-*r* reflects dissimilarities in the overall patterns of connectivity between the two maps, 1-eta^2^ also reflects differences in the connectivity strength for voxels with similar patterns of connectivity.

### Single-Feature Support Vector Machine Binary Classification of Diagnostic Status

We used support vector machine (SVM) binary classifiers to test whether individual cerebellar and striatal connectivity features could be used to distinguish between patients and controls. We masked each subject’s 13 individual seed-based connectivity maps with the significantly hyper-connected and hypo-connected regions from the main effect of group. We then calculated each subject’s average connectivity strength (Fz) within the hyper-connected mask and hypo-connected mask for each parcel’s functional connectivity map, resulting in 26 values per subject. The average hyper-connectivity and hypo-connectivity Fz for each network were then linearly combined, resulting in 13 values per subject, and each of these was used as a single feature to train an SVM (i.e. the 13 parcel connectivity features were used to train 13 separate classifiers; **Supplementary Fig. S34-S35**). We also trained a classifier on the average connectivity across all cerebellar parcels and on the average connectivity across all striatal parcels. Lastly, we trained classifiers using a linear combination of both the cerebellar and striatal parcel connectivity from each of the 6 networks (VIS was excluded as it had no striatal component) as well as the average connectivity of all 13 parcels (**Supplementary Fig. 36**). Hence there were a total of 13 individual parcel + 1 average cerebellum parcels + 1 average striatum parcels + 6 combined + 1 average all parcels = 22 single-feature classifiers. For each classifier, we ran 1,000 cross-validation runs. In each run, a subset of subjects (50%) was randomly selected and used to train the classifier, which was then tested on the remaining subset of subjects.

We employed receiver operating characteristic (ROC) curves to quantify the performance of the obtained classifiers. The ROC curve plots the true positive rate (TPR) against the false positive rate (FPR). TPR, or sensitivity, was calculated as the proportion of SCZ subjects correctly diagnosed as SCZ by the classifier. FPR, which is equal to (1–specificity), was calculated as the proportion of CON subjects incorrectly diagnosed as SCZ by the classifier. A “perfect” classifier that can accurately predict all new cases will thus have a point at (0,1) where TPR=1.00 (100% sensitivity) and FPR=0.00 (100% specificity), whilst a classifier operating at chance will fall along the 45° diagonal (“line of no discrimination”). We also calculated the Area Under the Curve (AUC), the area between the ROC curve and the X-axis, for all 1,000 runs of each classifier. The AUC is a summary statistic that can be interpreted as the probability that the classifier will accurately classify a randomly drawn pair of SCZ and CON subjects.

Finally, we performed hyper-parameter optimization of the SVM by repeating the process reported above using a range of parameters. Specifically, we tested soft margins (*C*) from 1.0e-5 to 1.0e5 at each order of magnitude. We also tested linear and nonlinear (Gaussian radial basis function with sigma=1) kernels. Additionally, we ran cross-validation using a 80%/20% split (as opposed to 50%/50%) between training and test samples. Notably, no other set of model parameters performed significantly better than the version reported here.

### Multi-feature SVM

To test if interactive effects between specific combinations of features contribute additional diagnostically relevant information, we used all 13 cerebellar and striatal parcel features to train one classifier in a multidimensional feature space (i.e. there were 13 values per subject). To perform feature selection, we used stepwise backward elimination, as follows: we ran 10,000 cross-validation runs on the 13-feature classifier and extracted the weights of each individual feature. We then removed the feature with the smallest absolute weight (i.e. the least discriminatory feature between SCZ and CON, here the striatal SOM); reran the 12-feature classifier; and calculated the mean AUC. We repeated this stepwise process, each time removing the least discriminatory network feature from the remaining features, until only one (here the striatal FPCN) was left. We also explicitly tested the interactive effect between the two features that were assigned the strongest weights in the multi-feature classifier, by adding the four-way interaction (striatum FPCN×cerebellum DAN; striatum DAN×cerebellum FPCN; striatum FPCN×striatum DAN; cerebellum FPCN×cerebellum DAN) as four additional features.

## RESULTS

### Bi-directional Brain-wide Cerebellar and Striatal Dysconnectivity is Observed in SCZ

Due to the length and scope of this study, a summary of our main hypotheses and findings, as well as the supporting figures and supplementary data, is provided in **Fig. 1**. First, to test whether SCZ is associated with uniform disturbances across CSTC networks, we quantified the functional connectivity of functionally distinct subdivisions in the cortex, thalamus, cerebellum and striatum, defined by a parcellation by Yeo et al. (2011), Buckner et al. (2011) and Choi et al. (2012) (Buckner RL *et al.* 2011; Choi EY *et al.* 2012). This parcellation, which was identified in 1,000 healthy adults and shown to be robust and replicable (Buckner RL *et al.* 2011; Yeo BT *et al.* 2011; Choi EY *et al.* 2012), contains both *sensory* networks (visual, VIS and somatomotor, SOM) and *associative* networks (dorsal attention, DAN; ventral attention, VAN; limbic, LIM; frontoparietal control, FPCN; and default mode, DMN) (**Supplementary Fig. S1**). Below, we use the term “parcel” to refer to a subdivision of the cortex, thalamus, striatum, or cerebellum belonging to one of the 7 functional networks. Of note, no representation of the visual network was discovered in the striatum (Choi EY *et al.* 2012); hence only 6 striatal parcels were used for all striatal analyses, and no representation of the limbic network was found in the thalamus. **Fig. 2a** shows the cortical (middle panel), cerebellar (left), and striatal (right) parcels of the 7 functional networks. Thalamic results are shown in **Supplementary Fig. S10**.

**Figure 1.**
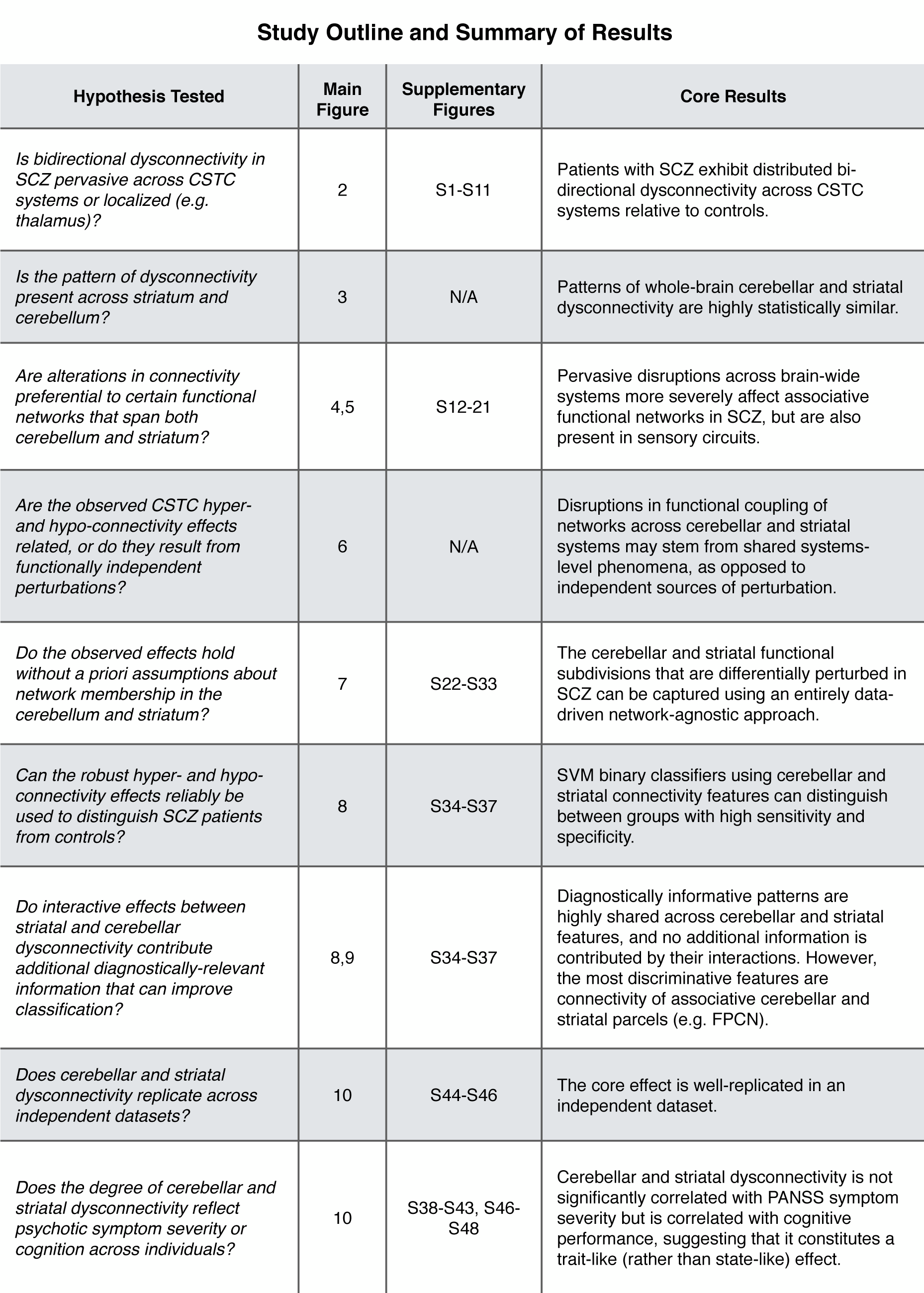
Summary of core questions and findings. Table of contents outlining the main questions studied in the present paper, the relevant figures in the main text and the supplemental materials, and the core results.

**Figure 2.**
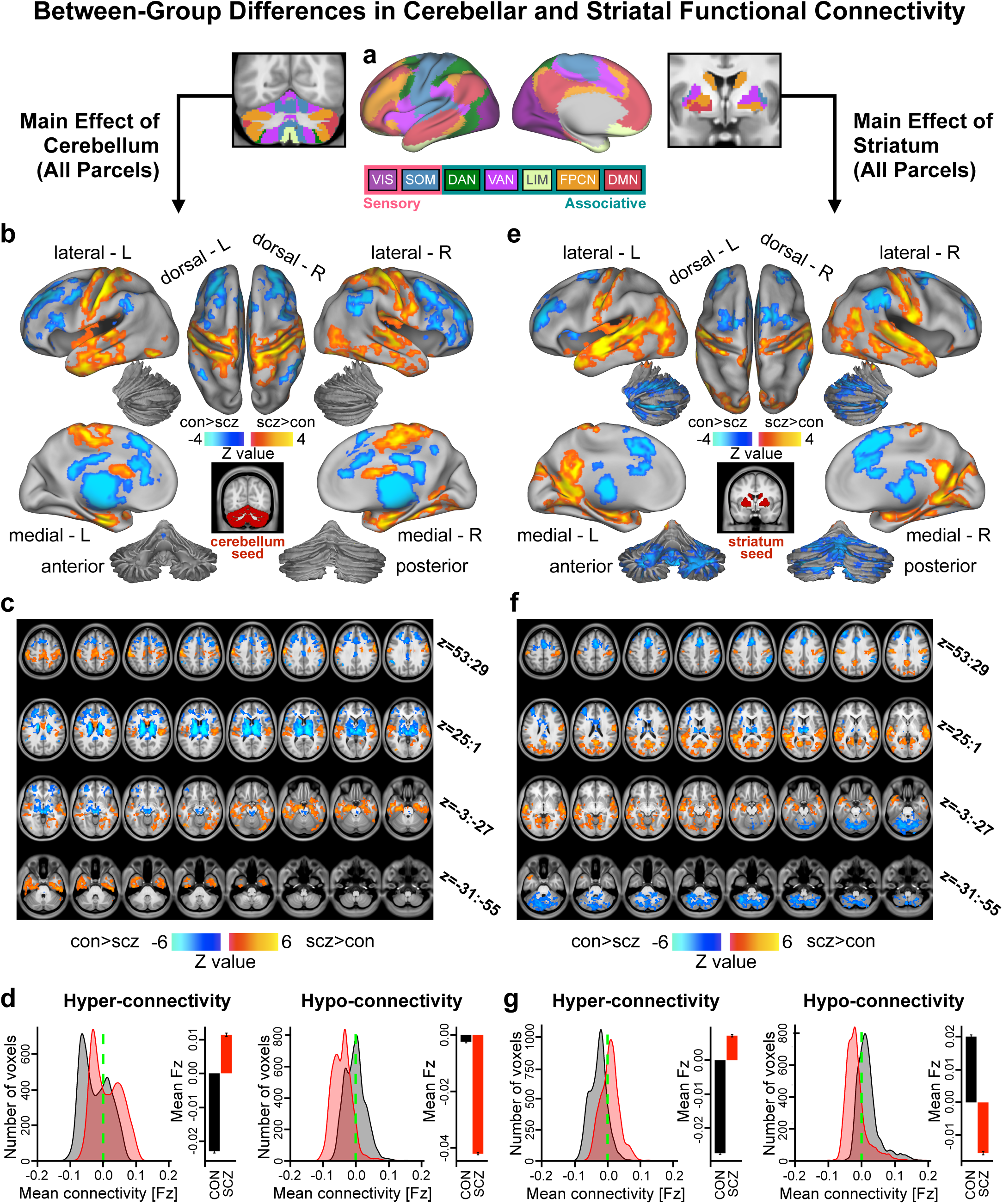
Main effect of cerebellar and striatal network parcel connectivity in SCZ. **(a)** Functionally defined networks obtained from 1,000 resting-state scans (first defined and independently replicated in 500 scans respectively) in the cerebral cortex, cerebellum and striatum (Buckner RL *et al.* 2011; Yeo BT *et al.* 2011; Choi EY *et al.* 2012). Whole-brain functional connectivity was computed from each of these 7 network parcels in the cerebellum (6 in striatum) and quantified in a *Group* (SCZ vs. CON group) x *Parcel* (VIS, SOM, DAN, VAN, LIM, FPCN and DMN) ANOVA. Networks were broadly grouped as either sensory or associative. **(b)** Cortical surface view of areas showing significant main effect of *Group* in whole-brain connectivity with the cerebellar parcels (*p<*0.05 TFCE whole-brain corrected) between 159 patients with chronic schizophrenia (SCZ) and 162 healthy controls (CON). Orange/yellow areas indicate regions where patients exhibited stronger cerebellar connectivity, whereas blue areas indicate regions where patients exhibited reduced cerebellar connectivity, relative to controls. Inset shows coverage of all cerebellar parcels. **(c)** Volume-based axial view of cerebellar connectivity group differences in **b** with Z-coordinate ranges (each slice in each row increments by 3 mm). **(d)** Distribution of connectivity strength (Fz) values within voxels showing significant hyper- and hypo-connectivity in SCZ and CON. Bar plots show mean connectivity averaged across all voxels in hyper- and hypo-connected areas. **(e-g)** Results for identical independent analysis conducted with the 6 functionally-defined striatal network parcels. Note that there was no representation of the visual network in striatum (Choi EY *et al.* 2012) and therefore that network was omitted. Abbreviations: VIS, visual network; SOM, somatosensory; DAN, dorsal attention network; VAN, ventral attention network; LIM, limbic network; FPCN, frontoparietal control network; DMN, default mode network.

We studied a well-powered sample of patients diagnosed with chronic SCZ (N=159) and matched healthy controls (CON, N=162; see **Table 1** for detailed demographics). First, we computed a 2×7 *Group* x *Parcel* ANOVA to test for differences in functional connectivity across the 7 cerebellar network parcels between SCZ and CON groups (see **Materials and Methods** and **Supplementary Fig. S2** for unthresholded Z-maps of all seeds). All reported whole-brain effects survived non-parametric threshold-free-cluster-enhancement (TFCE) correction (Smith SM and TE Nichols 2009) with 2,000 permutations (see **Materials and Methods**). Henceforth we use the term ‘dysconnectivity’ to denote the difference between SCZ and CON groups in the statistical covariation of BOLD signals over time across regions. Between-group differences revealed cerebellar hyper-connectivity with somatomotor cortex in SCZ relative to CON, and hypo-connectivity with prefrontal cortex, thalamus and striatum (**Fig. 2b-c**; see **Supplementary Fig. S4a-b** for unthresholded maps). **Fig. 2d** highlights the shift in the distribution of cerebellar connectivity strengths in SCZ for both hyper-connected and hypo-connected regions and the group difference in mean connectivity of these regions. An independent 2×6 *Group* x *Parcel* ANOVA was computed for the 6 functional striatal parcels (see **Supplementary Fig. S3** for unthresholded Z maps of all parcels). Between-group differences revealed qualitatively similar dysconnectivity patterns as with cerebellar parcels, namely hyper-connectivity with somatomotor cortex and hypo-connectivity with frontal cortex, thalamus, and cerebellum (**Fig. 2e-g;** see **Supplementary Fig. S4c-d** for unthresholded map). Repeating these analyses without global signal regression did not alter key effects (**Supplementary Fig. S5**). As expected, differences across parcels (i.e. main effect of *Parcel* irrespective of group) revealed widespread differential coupling for both cerebellar and striatal analyses, indicating that connectivity patterns differ across functional parcels (**Supplementary Fig. S6**). Critically, the reported between-group effects were not explained by smoking status, head motion, signal-to-noise ratio, or medication dose (**Supplementary Fig. S7**). Between-group effects adjusted for motion and signal-to-noise, as well as the effects of these covariates, are reported in **Supplementary Fig. S8-S9**.

To verify that the bi-directional pattern of dysconnectivity is in fact pervasive across CSTC systems, we conducted a similar analysis for functional connectivity of thalamic parcels (**Supplementary Fig. S10, Supplementary Materials and Methods**). These results replicate earlier findings of bi-directional thalamic connectivity in SCZ (including results previously reported in a subset of the same subjects used in present analyses) (Welsh RC *et al.* 2010; Woodward ND *et al.* 2012; Anticevic A, MW Cole*, et al.* 2014). Additionally, we studied the dysconnectivity of the entire CSTC functional pathway, simultaneously seeding from all parcels of the same network across cortex, striatum, thalamus, and cerebellum. Here we examined the effects of specific circuits, such as the CSTC-wide FPCN (**Supplementary Fig. S11**) because combining all networks across CSTC systems lacks specificity (i.e. it includes virtually all gray matter). As hypothesized, the bi-directional pattern of dysconnectivity was again observed, suggesting that these disturbances in functional relationships are indeed unified across neural systems. As noted, thalamic and cortical dysconnectivity in SCZ has been previously well-characterized (Welsh RC *et al.* 2010; Woodward ND *et al.* 2012; Anticevic A, MW Cole*, et al.* 2014; Baker JT et al. 2014) but never extended across CSTC. Therefore, we focus on the cerebellar and striatal components of the CSTC systems to examine similarity of disturbances.

Given the marked qualitative similarity between the cerebellar and striatal dysconnectivity maps (**Fig. 3a**, **Supplementary Fig. S4**), we formally quantified the overlap between the two independent *a priori* analyses. This was achieved by computing the correlation between the two group difference maps (expressed as difference in Fz values), but after removing cerebellar and striatal voxels from their respective maps. A significant positive relationship (*r*=0.40, *p*<0.001) indicated high similarly between striatal and cerebellar dysconnectivity in SCZ (**Fig. 3b**). We also quantified the cerebellar and striatal dysconnectivity overlap for areas showing significant hyper-connectivity (yellow) and hypo-connectivity (blue) (**Fig. 3c & d**). **Fig. 3e** indicates high overlap across both cerebellar and striatal hyper-connectivity effects (orange areas indicate overlapping effects, logical OR; yellow areas indicate joint effects, logical AND). Notably, the number of voxels in the overlapping areas was above chance (hypergeometric test for probability of overlap, *p<*0.001, given the total number of voxels in the brain) (**Fig. 3f**). Similarly, **Fig. 3g** indicates the same result for hypo-connectivity effects (dark blue areas indicate overlapping effects, logical OR; light blue areas indicate joint effects, logical AND). Again, the number of hypo-connected voxels that overlapped between cerebellar and striatal analyses was above chance (*p<*0.001, **Fig. 3h**). Overall, quantitatively and spatially high similarity between cerebellar and striatal dysconnectivity suggests highly comparable brain-wide perturbations.

**Figure 3.**
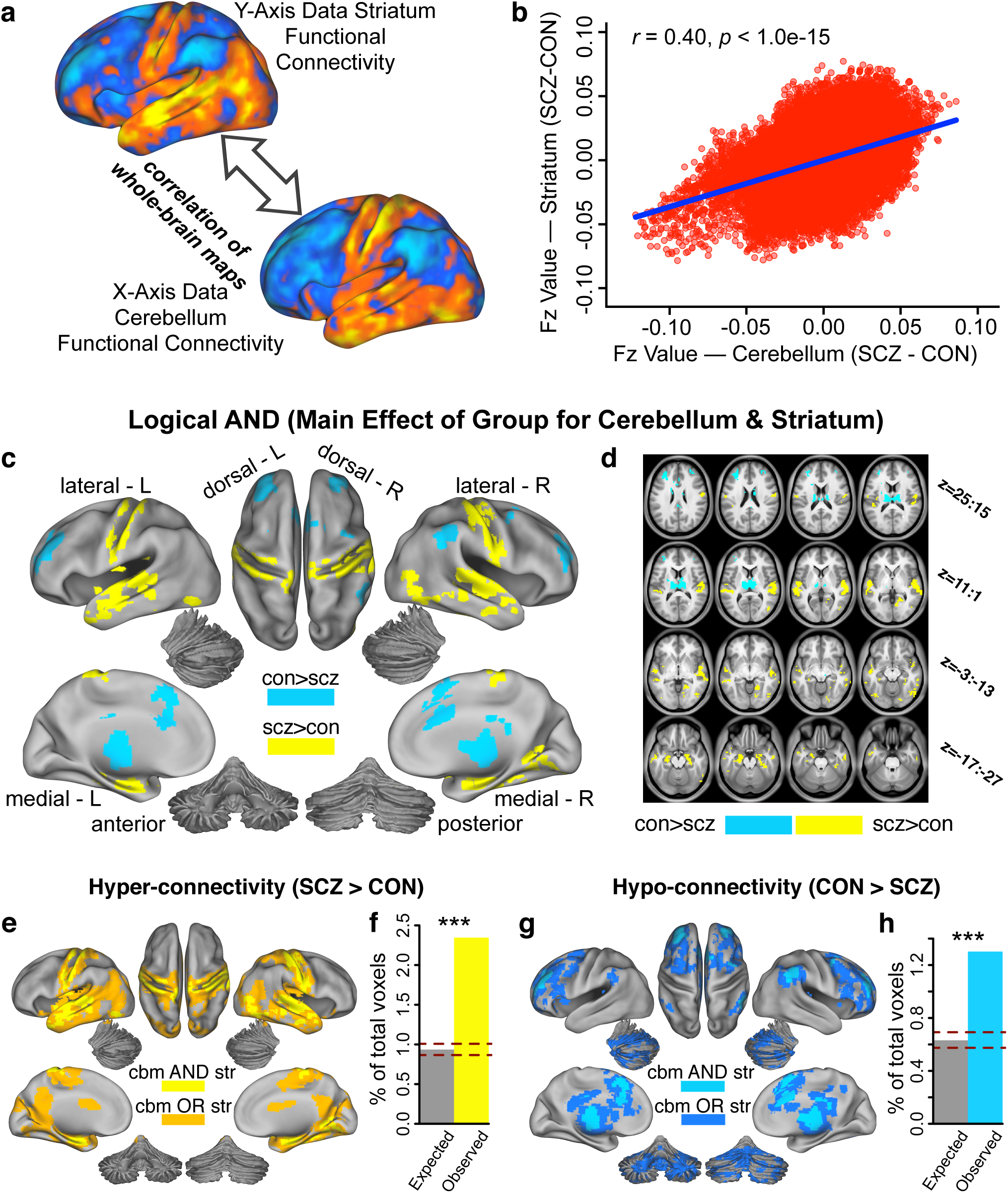
Similarity between perturbed cerebellar and striatal connectivity in SCZ. **(a)** Surface unthresholded Z-maps showing cerebellar and striatal between-group connectivity differences. Qualitatively, regions that exhibited an increase in cerebellar connectivity in SCZ also exhibited an increase in striatal connectivity, and vice versa. **(b)** Scatterplot showing a highly significant positive relationship between group differences in cerebellar and striatal whole-brain connectivity (*r*=0.40, *p*<1.00e-15). The plot shows Fz values for all voxels in cerebellar (X-axis) and striatal (Y-axis) between-group connectivity maps. **(c)** Surface maps of conjunction analysis showing areas that are significantly hyper-connected (yellow) and hypo-connected (blue) with both cerebellar and striatal parcels. **(d)** Volume-based axial view of **a** with Z-coordinate ranges (each slice in each row increments by 3 mm) **(e)** Surface view of areas showing both significant cerebellar (cbm) and striatal (str) hyper-connectivity (logical AND, bright yellow) and areas significantly hyper-connected to one or the other (logical OR, orange). **(f)** The number of voxels overlapping between observed cerebellar and striatal hyper-connectivity maps is significantly greater than chance (*p<*0.001), given the total number of voxels in the brain. **(g)** Surface view of areas showing both significant cerebellar (cbm) and striatal (str) hypo-connectivity (logical AND, bright blue) and areas significantly hypo-connected to one or the other (logical OR, dark blue). **(h)** Number of voxels that overlap between observed cerebellar and striatal hypo-connectivity maps is significantly greater than chance (*p<*0.001), given the total number of voxels in the brain. Dashed red lines in **f** and **h** indicate 95% Clopper-Pearson confidence intervals for chance.

### Dysconnectivity in SCZ is Driven by Associative Network Disruptions

Prior analyses are highly consistent with the hypothesis that CSTC disruptions in SCZ are robust and highly conserved across both the cerebellar and striatal systems. However, it remains unknown if these alterations are preferential to some functional networks that span both cerebellum and striatum. For instance, prior work strongly implicates associative network disturbances in SCZ in the cerebral cortex (Meyer-Lindenberg AS et al. 2005; Baker JT *et al.* 2014; Yang GJ, JD Murray, XJ Wang*, et al.* 2016). Yet, to our knowledge, no study has tested if such disruptions persist in a uniform pattern across the associative networks for both cerebellum and striatum. To test this we computed a *Group x Parcel* interaction, which would explicitly reveal differential between-group disruptions across distinct cerebellum and striatum parcels. The cerebellar *Group* x *Parcel* interaction effect revealed 37 areas throughout the brain (**Fig. 4a**). In turn, the striatal *Group* x *Parcel* interaction effect revealed 60 areas throughout the brain (**Fig. 4e**). These effects suggest that between-group dysconnectivity was not uniform across all parcels for either the cerebellar or striatal analyses. Importantly, these whole-brain interaction effects did not solely depend on within-cerebellum or within-striatum interactions (see **Supplementary Materials and Methods** and **Supplementary Fig. S12-S14**). Next, we tested for the source of the interaction and examined if the interaction effect was consistent for both cerebellum and striatum. Specifically, given the number of identified regions, we performed a hierarchical average-linkage clustering algorithm on the between-group connectivity of regions identified across the two interaction maps (**Materials and Methods**). For the cerebellum analysis, we identified three sets of areas that showed quantitatively similar patterns of dysconnectivity (**Fig. 4b**). The inset shows the distribution of areas in the three sets an axial slice (color-coded green, fuchsia and blue, as in the dendrogram). Next, we characterized the interaction effects for each set of areas produced by the clustering algorithm. Specifically, for a given striatal or cerebellar parcel/seed we averaged connectivity for all areas within a set produced by the clustering algorithm (**Fig. 4c**). We did this separately for both SCZ (red bars) and CON (black bars) groups. Each of the three sets of areas by definition shows quantitatively similar interaction effects (i.e. they are grouped by the clustering algorithm). The goal here is to characterize how the three sets of areas produced by the clustering algorithm differ. In Set 1, areas exhibited higher mean connectivity in SCZ compared to CON (i.e. hyper-connected on average). In Set 2, areas exhibited lower mean connectivity in SCZ (i.e. hypo-connected on average). The difference between SCZ and CON was less pronounced in Set 3 ROIs and exhibited a ‘mixed’ motif. Next, we averaged exclusively across sensory (VIS and SOM; pink) and associative (DAN, VAN, LIM, FPCN, DMN; teal) parcels to test if the observed alterations are preferential to higher-order networks. This analysis indicated that the between-group effect in each set is much more pronounced for associative parcels than sensory parcels (**Fig. 4d**). In essence, the entire source of the *Group* x *Parcels* interaction was driven by a more pronounced associative network alteration in the cerebellum.

**Figure 4.**
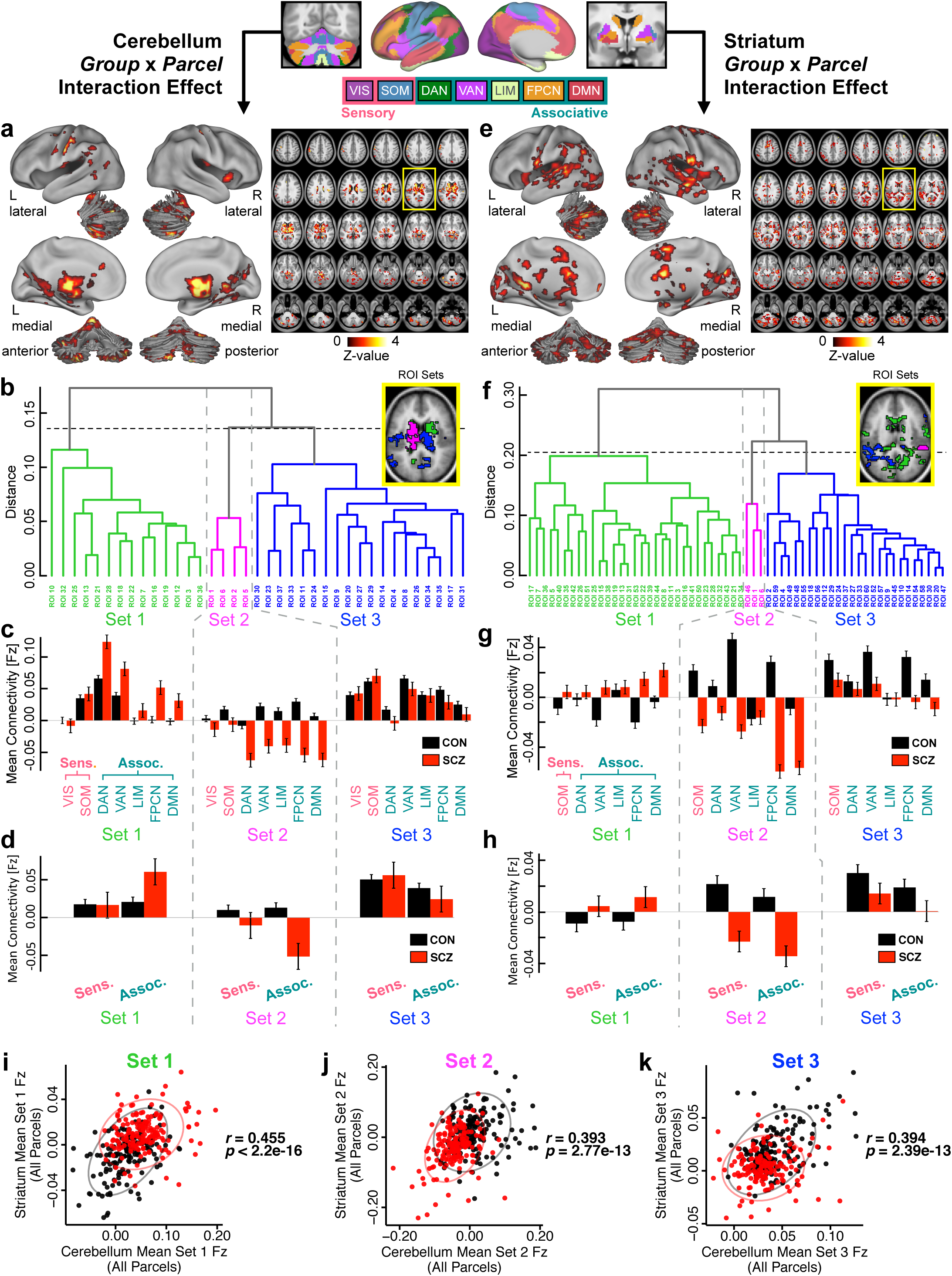
*Group* x *Parcel* interaction effect for cerebellar and striatal connectivity. Parallel analyses were conducted independently for the cerebellum and striatum, using the functionally-defined 7 cerebellar and 6 striatal parcels. **(a)** Surface view (left) and volume slice view (right) of regions of interest (ROI) revealing a significant *Group* x *Parcel* interaction effect for the cerebellar analysis. Slice outlined in yellow is magnified in inset in panel **b**. **(b)** Each of the 37 cerebellar interaction ROIs were assigned to one of three sets based on the between-group differences in functional connectivity of all 7 cerebellar functional parcels using an average-linkage hierarchical clustering algorithm, such that ROIs within a set share the most similar patterns of group differences in seed-based connectivity. Inset shows location of ROIs color-coded for each set. **(c)** To illustrate the sources of the interaction effect, the mean functional connectivity for each set of ROIs is shown separately for SCZ and CON subjects in each of the cerebellar parcels, averaged across all ROIs in the set. Error bars indicate standard error. ROIs in Set 1 exhibit higher mean connectivity in patients compared to controls, whereas ROIs in Set 2 exhibit lower mean connectivity in patients. **(d)** This effect is more pronounced for associative parcels (pink) than for sensory parcels (teal). ROIs in Set 3 are less differentiated between patients and controls. **(e-h)** Parallel analysis conducted for the 60 interaction ROIs from 6 striatal functional parcels. **(i-k)** Correlations between mean functional connectivity (Fz) of all cerebellar and striatal parcels in **(i)** Set 1, **(j)** Set 2, and **(k)** Set 3 interaction ROIs, across SCZ and CON subjects. Displayed *r* values are across all subjects (N=320, one outlier removed). Abbreviations: Sens., sensory networks; Assoc., associative networks; VIS, visual network; SOM, somatomotor network; DAN, dorsal attention network; VAN ventral attention network; LIM, limbic network; FPCN, frontoparietal control network; DMN, default mode network.

Next, we repeated the analyses for the striatal interaction effects. Strikingly, as with the cerebellum, a hierarchical clustering algorithm revealed three sets of areas with similar patterns of connectivity (**Fig. 4f**). The group differences in connectivity in each of these sets driving the striatal interaction effect was similar to those in the cerebellar interaction effect – namely, higher mean connectivity in SCZ than CON for ROIs in Set 1, lower mean connectivity in SCZ than CON for ROIs in Set 2, and a less ostensible difference for ROIs in Set 3 (**Fig. 4g**). These between-group differences were not as pronounced in striatal associative versus sensory parcels than in the analogous cerebellar analysis (**Fig. 4h**) because there was only one sensory parcel in the striatum. Finally, we computed the statistical similarity of these sets of ROIs between the cerebellum and striatum (**Fig. 4i-k**). Specifically, for each subject, we averaged Fz values across all ROIs in each cerebellar and striatal set. We did so for each parcel within the striatum and the cerebellum (e.g. the FPCN parcel). This revealed a robust and consistent relationship between cerebellar and striatal values across all sets. This strongly supports consistency of interactive effects for cerebellar and striatal ROIs for individual subjects. This relationship held for all networks (Set 1: *r*=0.455, *p*<2.20e-16; Set 2: *r*=0.393, *p*=2.77e-13; Set 3: *r*=0.394, *p*=2.39e-13) and particularly for the FPCN (Set 1: *r*=0.538, *p*<2.20e-16; Set 2: *r*=0.475, *p*<2.20e16; Set 3: *r*=0.367, *p*=1.16e-11; see **Supplementary Fig. S15** for all individual parcels/networks). Collectively, these analyses suggest that interactive brain-wide network disturbances are highly similar between the cerebellum and striatum. However, they are particularly obvious and highly consistent in certain functional networks, especially those involved in higher-order executive tasks.

Next, to further establish that the SCZ dysconnectivity was more pronounced in associative than in sensory networks, we examined seed-based functional connectivity maps of each parcel. Qualitatively, the between-group dysconnectivity for associative parcels closely matched the main group dysconnectivity effect (i.e. across all parcels, **Fig. 2b** and **2e**). This was markedly less prominent for sensory parcels across both cerebellum and striatum. To quantitatively verify this, we examined the similarity of the FPCN parcel (**Fig. 5a-b, g-h**) and the SOM parcel between-group effects (**Fig. 5d-e, j-k**) in relation to the main effects across both cerebellum and striatum. Specifically, we correlated unthresholded whole-brain between-group maps for the FPCN parcel with the unthresholded overall dysconnectivity maps (i.e. maps in **Fig. 2b** and **2e**). **Fig. 5c** highlights the similarity between FPCN and overall effects across all parcels (Cerebellum: *r*=0.914, *p*<1.0e-15; Striatum: *r*=0.913, *p*<1.0e-15). In contrast, the SOM effect dysconnectivity was substantially less similar to the main effects for the cerebellum and striatum (*r*=0.696, *p*<1.0e-15; *r*=0.754, *p*<1.0e-15). This constituted a highly significant difference between correlations for the FPCN and SOM similarity (Cerebellum: *t*=-255.12, *p*<1.0e-15, Striatum: *t*=-124.43, *p*<1.0e-15, William’s test for dependent correlations). The full expansion of all pair-wise thresholded and unthresholded maps is shown in **Supplementary Fig. S16-S21.** Collectively, these effects buttress prior analyses by showing pervasive disruptions across brain-wide networks that preferentially affect associative functional systems in SCZ.

**Figure 5.**
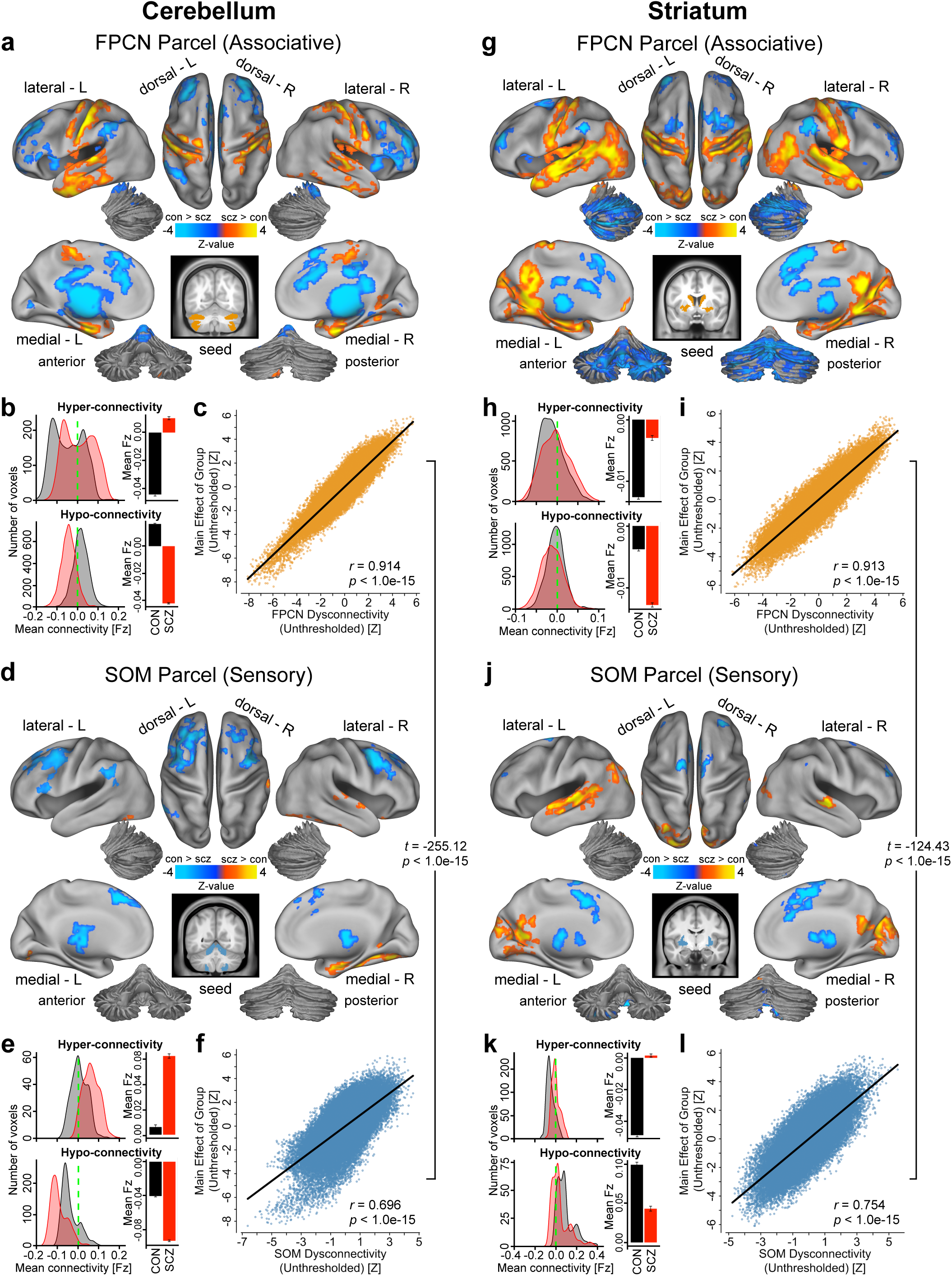
Group connectivity differences for associative versus sensory cerebellar and striatal parcels. **(a)** Surface view of regions for which connectivity with the cerebellar FPCN parcel, an exemplar “associative” parcel, showed significant between-group differences, *p*<0.05. **(b)** Density plots show distribution of connectivity strength (Fz) values within voxels showing significant hyper- and hypo-connectivity. Bar plots show mean connectivity averaged across all voxels in hyper- and hypo-connected areas. **(c)** Correlation between the voxelwise unthresholded whole-brain group dysconnectivity maps from the cerebellar FPCN parcel and the cerebellar main effect of group. Similar analyses are shown for the **(d-f)** cerebellar SOM parcel, **(g-i)** striatal FPCN parcel, and **(j-l)** striatal SOM parcel. Note the high resemblance between the associative parcel effect and the main effect of group for both the cerebellar and striatal analyses. Contrarily, the sensory parcel effects are significantly less similar (Cerebellum: *t*=-255.12, *p*<1.0e-15, Striatum: *t*=-124.43, *p*<1.0e-15, William’s test for dependent correlations). Abbreviations: FPCN, frontoparietal control network; SOM; somatosensory network.

### Relationship Between Hyper-connectivity and Hypo-connectivity Across Individuals

Presented analyses highlight that the alterations are strongly driven by associative networks within the cerebellum and striatum. However, more broadly, it is unknown if the observed hyper-connectivity and hypo-connectivity effects that dominate all presented analyses arise are linked. Alternatively, the bi-directional disruption may result from functionally independent perturbations: distinct groups of subjects could be driving the hyper-vs. hypo-effects across all analyses. To arbitrate between these possibilities, we quantified the connectivity strengths across subjects for areas showing hyper- and hypo-connectivity generally. There was a significant negative relationship between cerebellar hypo-and hyper-connectivity in CON (*r*=-0.77, *p*<2.2e-16), indicating that control subjects with the weakest coupling between cerebellar-sensorimotor regions also show the strongest coupling between cerebellar-thalamo-striatal-prefrontal regions (**Fig. 6a**). This effect replicated in the SCZ sample (*r*=-0.49, *p*<4.1e-11), but it was significantly attenuated compared to CON (*Z*=-4.29, *p*<0.001). Similarly, CON subjects with the weakest striatal-sensorimotor coupling showed the strongest striatal-thalamo-prefrontal-cerebellar coupling (*r*=-0.85, *p*<2.2e-16), and an attenuated effect was seen in SCZ subjects (*r*=-0.67, *p*<2.2e-16; *Z*=-3.95, *p*<0.001; **Fig. 6b**). Additionally, CON subjects that showed the strongest coupling with cerebellum also exhibited the strongest coupling with striatum, for both the hyper-connectivity (*r*=0.64, *p*<2.2e-16, **Fig. 6c**) and hypo-connectivity (*r*=0.75, *p*<2.2e-16; **Fig. 6d**) effects. This relationship was also present but attenuated in SCZ subjects (hyper-connectivity: *r*=0.34, *p*<1.5e-5, *Z*=3.58, *p*<0.001; hypo-connectivity: *r*=0.60, *p*<2.2e-16, *Z*=2.48, *p*=0.013). Note that in all **Fig. 6** panels, SCZ subjects show a diagonal ‘shift’ relative to CON across both zero-lines (dashed green), suggesting that the differences in hyper- and hypo-connectivity in SCZ may share a source of disturbance. Collectively, these analyses support the hypothesis that perturbations in information flow across these networks may stem from related systems-level phenomena, as opposed to functionally independent perturbations.

**Figure 6.**
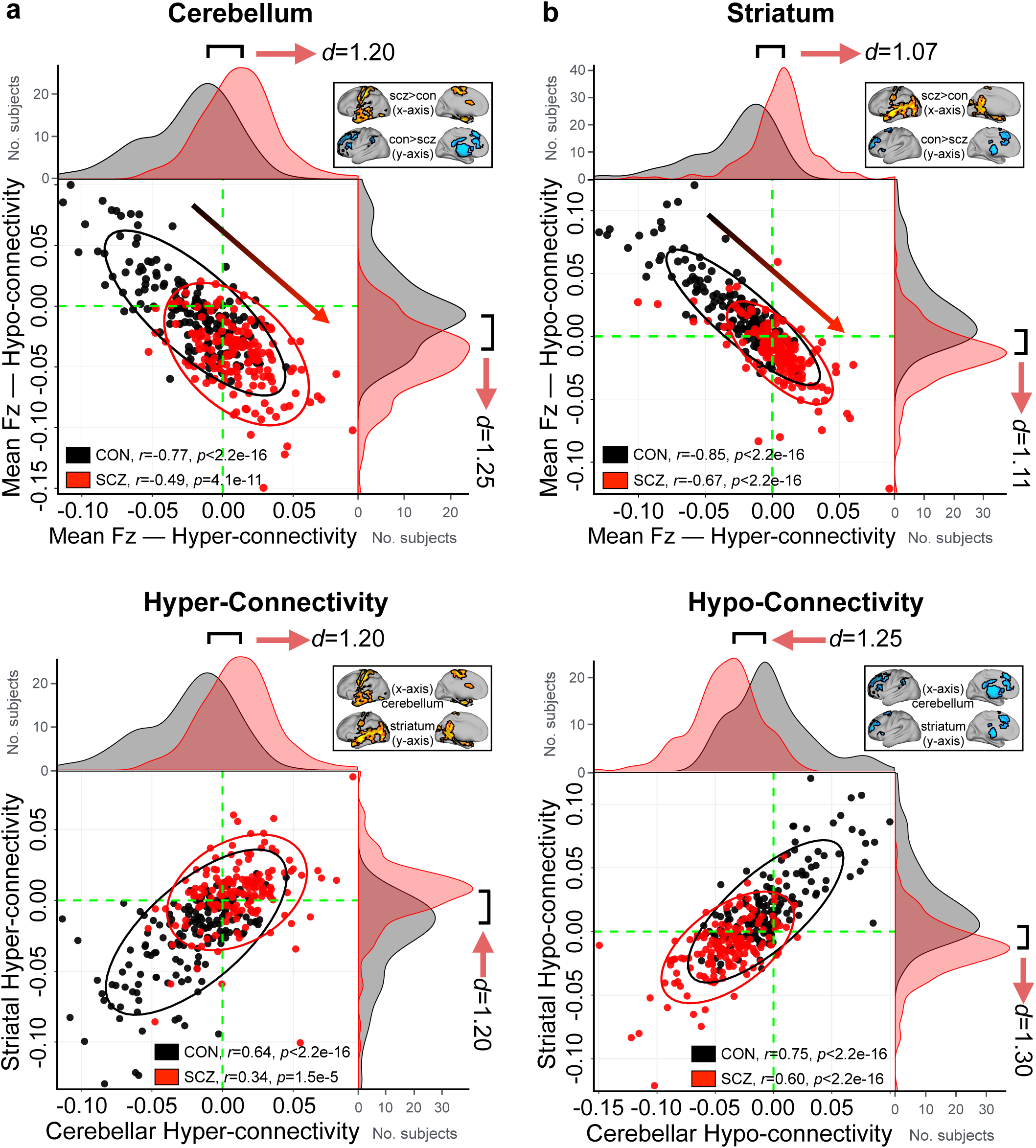
Relationship between cerebellar and striatal hyper- and hypo-connectivity across subjects. (a) Significant negative relationship evident between average hyper- and hypo-whole-brain connectivity with the cerebellum across all CON (black data points). SCZ (red data points) showed a ‘shift’ across the zero lines, indicating weaker prefrontal-striatal-thalamic-cerebellar coupling but stronger somatomotor-cerebellar coupling. Ellipses for each group mark the 95% confidence intervals. Distributions of average connection strengths for each subject show a shift in cerebellar coupling in SCZ, highlighting increased cerebellar coupling with somatomotor regions and decreased coupling with prefrontal-striatal and thalamic regions for patients. Inset shows regions of hyper- and hypo-connectivity with the cerebellum in SCZ, from **Fig. 2b. (b)** A similar effect is seen in an independently conducted analysis on whole-brain striatal connectivity. Again, SCZ showed a shift across the zero lines relative to CON, suggesting that the differences in hyper- and hypo-connectivity observed in SCZ may share a source of disturbance. Inset shows regions of hyper- and hypo-connectivity with the striatum in SCZ, from **Fig. 2e**. **(c)** CON subjects with the highest cerebellar connectivity in hyper-connected regions also show highest connectivity in striatal hyper-connected regions. Shift in SCZ subjects relative to CON is evident for both cerebellar and striatal hyper-connectivity (identical data as those plotted along X-axes in panels **a** and **b**), suggesting the underlying disruption is linked across these two systems. **(d)** Similarly, SCZ subjects show a shift in both cerebellar and striatal hypo-connectivity.

### Data-Driven Clustering of Cerebellar and Striatal Dysconnectivity

While robust, all presented analyses have used *a priori* functional networks to reveal shared CSTC perturbations in SCZ. However, there may be important functional differences in the spatial distribution of these network parcels in patients. In other words, patients may not show a clean separation that is assumed to be present in controls of cerebellum and striatum networks. Therefore, to test if the effects hold without *a priori* network assumptions, we performed data-driven *k-*means clustering on voxelwise group dysconnectivity for both the cerebellum and the striatum (see **Materials and Methods** and **Supplementary Fig. S22** for details). If the hypothesized *a priori* disruptions are indeed intrinsic and not attributable to spatial network assumptions, then the results should replicate via data-driven between-group parcellation of the cerebellum and striatum. We report several cluster solutions of between-group differences (see **Supplementary Fig. S23** for full 4-, 6- and 7-cluster solutions). While the pattern is highly similar irrespective of cluster choices, here we highlight the 7-cluster solution for the cerebellum and the 6-cluster solution for the striatum to allow a direct comparison with the *a priori* network parcellation granularity (i.e. this was the number of networks in the *a priori* parcellations).

The clustering solution for the cerebellum revealed well-defined and bilaterally symmetrical clusters, which is not consistent with the effects being driven by artifact (**Fig. 7a**). A voxelwise measure of dissimilarity (1-eta^2^; see **Materials and Methods**) shows regions with the numerically greatest degree of dysconnectivity between SCZ and CON. These foci were also bilaterally symmetrical (**Fig. 7b**). Next, we tested if these data-driven clustering solutions map onto the *a priori* network parcel results. Here we quantified the relationship between the group dysconnectivity for each of the obtained clusters and each *a priori* cerebellar parcel (top 7 rows) as well as the average across all parcels (bottom row). This yields a 8×7 matrix of Pearson’s *r* values shown in **Fig. 7c**, where the columns of the matrix are ordered such that the trace of the parcel-by-cluster square matrix is maximized. As illustrated by the diagonal, there was a selective mapping between data-driven and *a priori* dysconnectivity patterns. Notably, six of the 7 clusters have their strongest correlation value along the diagonal, suggesting that these clusters mapped selectively to existing parcels. This selective mapping can also be seen with the alternative *k*=4 and *k*=6 cluster solutions (**Supplementary Fig. S32**). Collectively, this indicates qualitatively similar disruptions irrespective of analytic methods used to define cerebellar dysconnectivity in SCZ. This is corroborated, for instance, by examining Cluster 4 (**Fig. 7a**, bright green), which is most similar in dysconnectivity to the associative parcels (*r=*0.81, *p<*0.001, **Fig. 7d**), indicating that in the context of SCZ-related disturbances it exhibits a functionally ‘associative-like’ profile. We show the threshold-free between-group differences in whole-brain connectivity of this ‘associative-like’ cerebellar cluster in **Fig. 7e** (see **Supplementary Fig. S24-S26** for maps of all clusters for 4, 6, and 7-cluster solutions).

**Figure 7.**
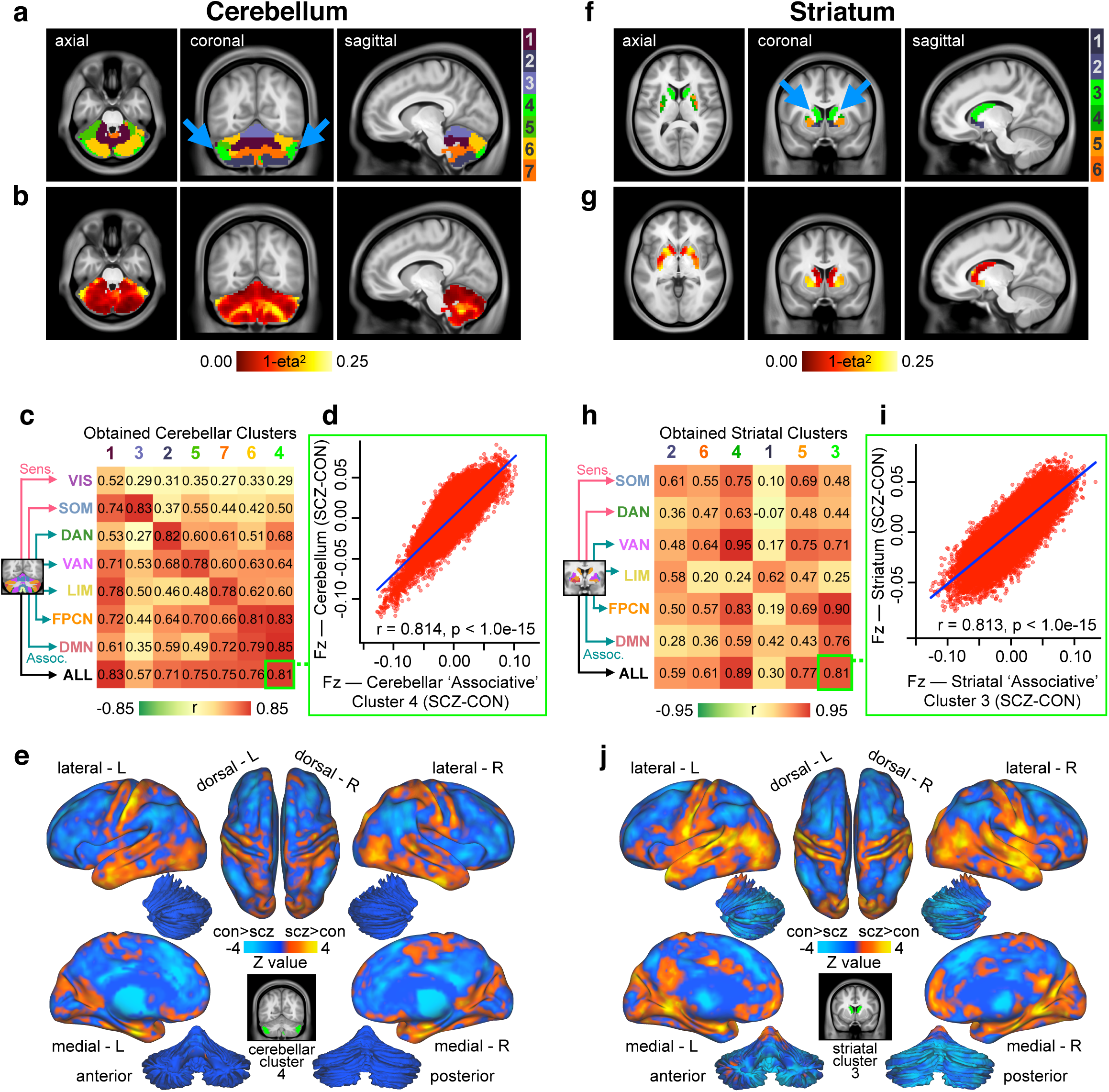
Voxelwise clustering of cerebellar and striatal group difference in connectivity. **(a)** Results for *k-*means 7-cluster solution identifying cerebellar voxels with the most similar patterns of between-group connectivity differences. Note the cluster solution is highly symmetrical (blue arrows). **(b)** Cerebellar dysconnectivity based on group dissimilarity. Brightest voxels indicate greatest between-group differences. **(c)** Matrix of correlations (Pearson’s *r*) between dysconnectivity of obtained clusters and dysconnectivity of functional parcels. Cluster numbers (column headings) are colored as in **a**. Functional parcels (row headings) are labeled and colored as in **Figure 2a** and original parcellation (Yeo BT *et al.* 2011). Columns have been ordered such that the trace of the parcel-by-cluster square matrix is maximal among all permutations. Cluster-seeded dysconnectivity shows a range of correlation strengths with functional parcel-seeded dysconnectivity. Cluster 4 (bright green) is strongly correlated with associative parcels, as well as with the overall main effect of group, suggesting that it is a functionally ‘associative-like’ cluster. **(d)** Scatterplot showing high degree of correlation between voxelwise whole-brain group dysconnectivity (Fz values) of cerebellar cluster 4 (the ‘associative’ cluster) and voxelwise whole-brain group dysconnectivity (Fz values) across all cerebellum parcels. **(e)** Surface unthresholded Z-maps showing between-group connectivity for cerebellar executive cluster (shown in inset). **(f-j)** Identical analyses conducted independently for between-group differences in whole-brain striatal connectivity. Striatal cluster 3 (bright green) is strongly correlated with associative parcels, as well as with the overall main effect of group, suggesting that it is a functionally ‘associative-like’ cluster.

Importantly, *k-*means clustering with *k=*6 on striatal voxelwise dysconnectivity yielded a consistent and bilaterally symmetrical solution (**Fig. 7f**). The most dissimilar voxels are again indicated in bright yellow in **Fig. 7g,** suggesting a non-uniform pattern of striatal alterations. As with the cerebellum, there was a selective mapping between data-driven and *a priori* dysconnectivity patterns for the striatum (**Fig. 7h**). Cluster 3 (**Fig. 7f**, bright green) exhibited a pattern of dysconnectivity most similar to those of associative functional parcels (*r=*0.81, *p<*0.001, **Fig. 7i**). Finally, threshold-free between-group differences in whole-brain connectivity of this ‘associative-like’ striatal cluster highlight a pattern that in line with the *a priori* association parcel analysis (**Fig. 7j**, see **Supplementary Fig. S27-S29** for maps of all clusters for 4, 6, and 7-cluster solutions). Additionally, we report the same data-driven clustering solutions for cerebellum and striatum in CON only (**Supplementary Fig. S33**), which fully replicated the functional parcels identified previously by Bucker et al. and Choi et al. (Buckner RL *et al.* 2011; Choi EY *et al.* 2012).

### Cerebellar and Striatal Dysconnectivity Features Predict Diagnostic Group Status

Above, we show consistent and robust alterations in BOLD functional relationships across striatal and cerebellar functional subdivisions in SCZ. Next, we examined if these effects may yield a sensitive and specific binary classification of diagnostic group status. Specifically, we used a support vector machine (SVM) binary classifier to test if cerebellar and striatal parcel connectivity features reliably distinguish between groups. In turn, we tested three secondary questions related to the brain-wide patterns of disruptions: First, we tested if any of the identified features selectively ‘drive’ the classification performance or if the classifier performance remains stable irrespective of parcel feature used. Put differently, if SCZ is associated with brain-wide alterations across CSTC systems, then cerebellar and striatal parcel features should yield comparable classification performance, as there should be overlap in the disruptions. Second, a corollary of this hypothesis is that specific parcels within the striatum or cerebellum may drive classifier performance. In particular, given that our previous analyses show preferential SCZ disruptions in associative parcels, we hypothesized that associative parcel features would yield the highest classification performance. However, if the brain-wide disruptions are shared across CSTC systems, then the same network parcels should contribute information to the classifier irrespective of whether they are selected from the striatum or the cerebellum. Third, we examined whether classifier performance could be improved by combining select subsets of cerebellar and striatal parcel features. We hypothesized that if there exists considerable overlap in diagnostically relevant information across cerebellar and striatal parcels, a classifier using a combination of all parcel features would not markedly outperform the most discriminatory single-feature classifier. Alternatively, a multi-feature classifier would yield better performance if interactive effects across specific cerebellar and striatal parcel features contribute unique diagnostically relevant variance.

First, to test if cerebellar and striatal features are comparable for classification, we used the connectivity strengths (Fz values) of hyper- and hypo-connected areas from each of the seed-based functional connectivity analyses as input features. Each classifier was trained on a randomly selected subset (50%) of subjects and tested on the remaining subjects, repeated 1,000 times (cross-validation runs, see **Materials and Methods**). Classifier performance across specific exemplar parcel features is shown in **Fig. 8** (see **Supplementary Fig. S34-S36** for all classifiers). Cerebellar and striatal classifiers yielded highly comparable performance, supporting the hypothesis of shared brain-wide disruptions across both the cerebellum and striatum. Specifically, the classifier trained on the average dysconnectivity across all parcels (AVERAGE) was similar for the cerebellum (accuracy=75.0%, sensitivity=76.9% & specificity=73.2%, mean area under the receiver operating characteristic curve (AUC)=0.835, **Fig. 8a**) and the striatum (accuracy=75.7%, sensitivity=79.0% & specificity=72.5%, AUC=0.853, **Fig. 8d**). Collectively, these measures suggest that the cerebellum and striatum may exhibit shared disruptions. Consequently, cerebellum and striatum do not outperform one another.

**Figure 8.**
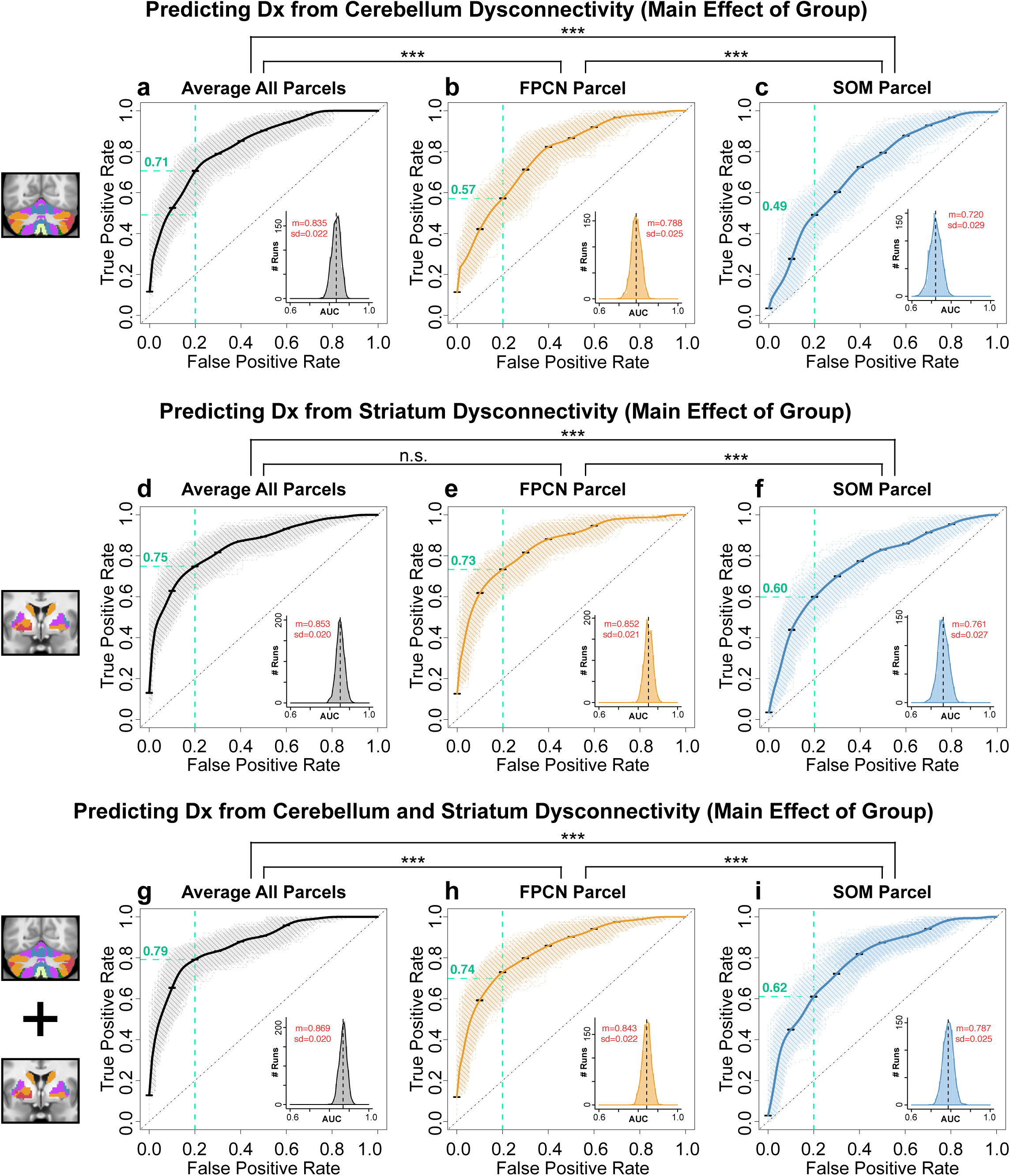
Receiver operating characteristic (ROC) curves from binary classifiers of diagnostic status trained on cerebellar and striatal dysconnectivity features. Exemplar associative (FPCN) and sensory (SOM) parcels, as well as the average combined across all parcels (AVERAGE), are shown. **(a)** Performance of classifier using AVERAGE dysconnectivity across all cerebellar parcels. Bold black curve plots the mean true positive rate (TPR) at each false positive rate (FPR) across 1,000 cross-validation runs (standard error bars shown in black). Grey curves show individual results from all cross-validation runs. Inset plots the distribution of the Area Under the Curve (AUC) for all 1,000 runs; black dashed line shows the mean AUC. Mean (m) and standard deviation (sd) of the AUC distribution are displayed in red. **(b)** Performance of classifier using cerebellar FPCN dysconnectivity. **(c)** Performance of classifier trained using cerebellar SOM parcel dysconnectivity. Green dashed lines show the TPR for each classifier at FPR=0.20. The cerebellar FPCN TPR=0.71, indicating that this classifier achieves a higher sensitivity than the SOM classifier (TPR=0.49) at the same ‘cost’ of specificity. Similarly, the AUC performance is significantly higher for the FPCN than the SOM classifier (*t*=55.692, df=1953.4, *p*<1.0e-15). Asterisks denote significance at *p*<0.001 in a two-tailed t-test between the indicated AUC distributions. **(d-f)** Performance of classifiers using striatal AVERAGE, SOM, and FPCN features. Again, the FPCN classifier outperformed the SOM classifier, as indicated by sensitivity at FPR=0.20 (FPCN TPR=0.73, SOM TPR=0.60) and AUC (*t*=83.869, df=1871.2, *p*<1.0e-15). **(g-i)** Performance of classifiers using combined cerebellar and striatal AVERAGE, FPCN, and SOM features. See **Supplementary Fig. S34**-**S36** for ROC curves of all parcels.

As noted, our second classifier hypothesis was to test if disruptions in SCZ are more pronounced in associative parcels relative to other parcels within cerebellar and striatal systems. This would result in better performance for classifiers trained on associative parcel features relative to sensory parcel features. Both the cerebellar and striatal FPCN classifiers outperformed the SOM classifiers (Cerebellum-FPCN Classifier: accuracy=71.2%, sensitivity=80.6% & specificity=61.8%, AUC=0.788, **Fig. 8b**; Striatum-FPCN Classifier: accuracy=75.7%, sensitivity=71.3% & specificity=71.0%, AUC=0.852, **Fig. 8e**; Cerebellum-SOM Classifier: accuracy=66.1%, sensitivity=76.9% & specificity=61.0%, AUC=0.720, **Fig. 8c**; Striatum-SOM Classifier: accuracy=68.3%, sensitivity=77.1% & specificity=59.7%, AUC=0.761, **Fig. 8f**). This was supported by a formal test of differences in FPCN vs. SOM AUC parameters across the cerebellum (*t*=55.692, df=1953.4, *p*<1.0e-15) and striatum (*t*=83.869, df=1871.2, *p*<1.0e-15). This result is consistent with the hypothesis of shared CSTC disruptions being preferential to associative parcels in SCZ.

In turn, our third classifier hypothesis tested if combining cerebellar and striatal information would improve classifier performance relative to the single best feature. We first trained classifiers on a linear combination of both cerebellar and striatal parcel features (referred to as “combined” classifiers below). The combined AVERAGE classifier yielded the highest performance (mean accuracy of 78.7%, with 80.7% sensitivity and 76.7% specificity, AUC=0.869, **Fig. 8g**). This was comparable to the combined FPCN classifier (accuracy=74.2%, sensitivity=80.8%, specificity=67.6%, AUC=0.843, **Fig. 8h**). Again, the combined classifier trained with SOM yielded the lowest performance (accuracy=70.8%, sensitivity=75.7%, specificity=66.0%, AUC=0.787, **Fig. 8i**). We then examined if interactive effects between these striatal and cerebellar parcel features contribute information that improves classification. Here we trained a 13-feature classifier using connectivity from all 7 cerebellar and 6 striatal parcels as predictors. This multi-feature classifier did not perform better than the classifiers trained on one parcel feature per subject (accuracy=77.3%, sensitivity=78.6% & specificity=76.0%, AUC=0.859, **Fig. 9a**), suggesting that interactions between parcel features did not contribute unique additional information. In further support of this effect, no added number of additional features appreciably improved classifier performance compared to the classifier trained on the single most discriminatory feature (**Fig. 9b**). Additionally, the relationship between the AUC of individual features and the weight assigned to them in the multi-feature classifier was not significant, suggesting that no informational tradeoffs were made by selecting certain parcel features over others in the 13 multi-feature classifier (**Fig. 9c**; all features: *r*=0.420, *p*=0.153, dashed grey line; excluding most discriminatory feature: *r*=0.11, *p*=0.74, blue line). Lastly, classifier performance was not improved by including the interactions between the two most strongly weighted parcel features – namely cerebellar and striatal FPCN and DAN – as additional predictors (accuracy=77.1%, sensitivity=79.1% & specificity=75.1%, AUC=0.856, **Fig. 9d**). Importantly, the classifier was not driven by group differences in head motion or SNR (**Supplementary Fig. 37**). These analyses suggest that diagnostically informative patterns are highly shared across cerebellar and striatal features, and no additional information is contributed by their interactions across these analyses.

**Figure 9.**
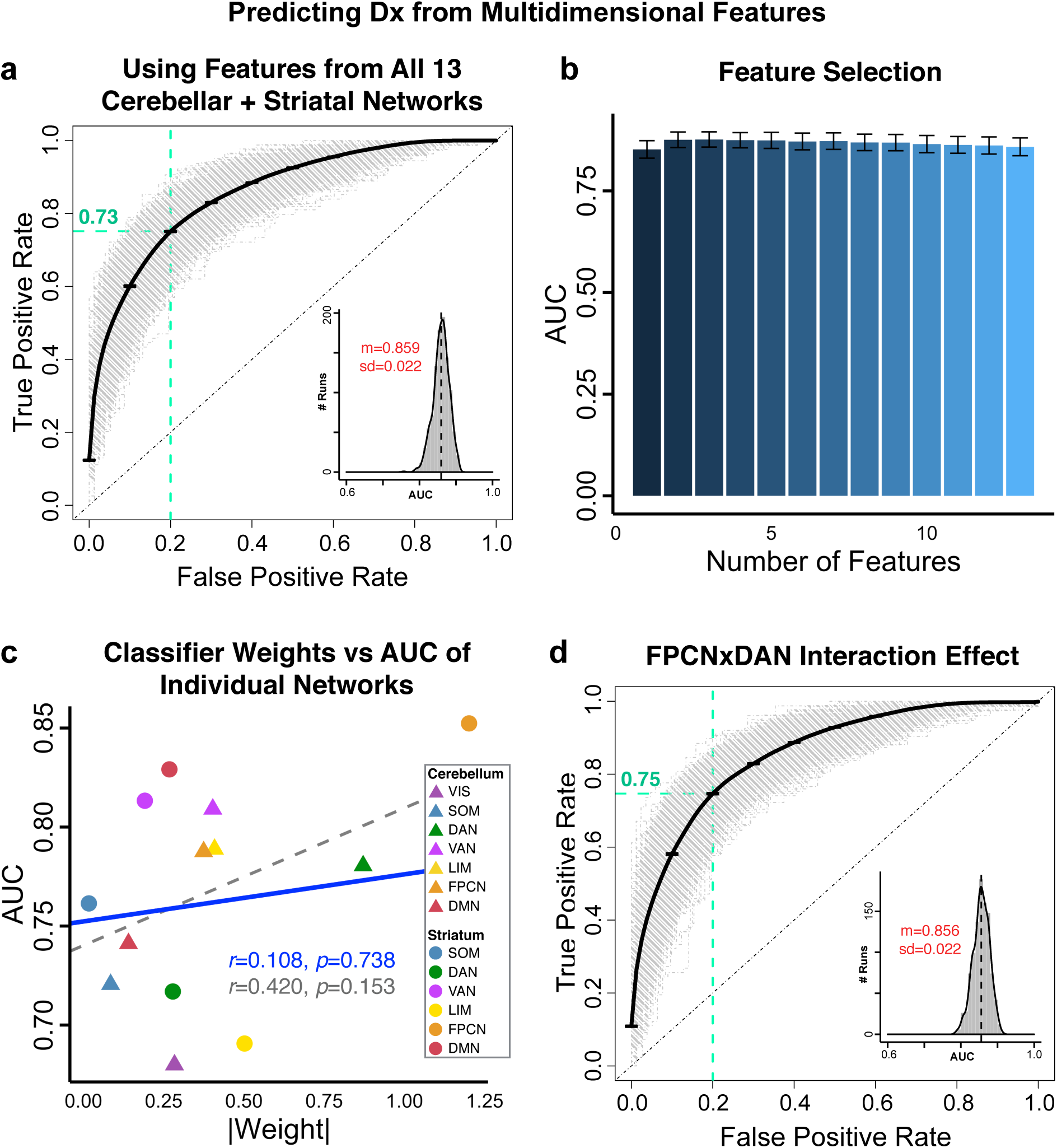
Binary classifiers using multidimensional predictors. **(a)** We trained a linear kernel SVM to distinguish between SCZ and CON subjects using connectivity features from all 7 cerebellar and 6 striatal parcels, in a 13-dimensional feature space. This multi-feature classifier did not perform significantly better than classifiers trained on individual features (**Fig. 8**, **Supplementary Fig. S34-S36**). ROC curves show the TPR plotted against the FPR of the classifier trained on all 13 parcel features. Grey curves show ROC curves for 1,000 individual cross-validation runs; bold black curve shows mean ROC curve (standard error bars in black). Green dashed lines indicate TPR=0.73 at FPR=0.20. Inset shows the distribution of the AUC across all 1,000 runs (black dashed line shows mean), with mean AUC (m) and standard deviation (sd) displayed in red. **(b)** Comparison of AUC performance across classifiers with 1 through 13 features. There were no significance differences in AUC between any of these classifiers, indicating that no additional number of features improved performance compared to the classifier trained on the single most discriminatory feature. Bar plots show the mean AUC of each classifier; error bars show standard deviation. **(c)** Plotting the mean AUC from classifiers trained on each individual parcel feature (from **Supplementary Fig. S34-S35**) against the absolute weight of these parcels in the 13-feature classifier shows that there is no linear dependence between the two. Dashed grey line shows correlation between all points (*r*=0.42, *p*=0.15); solid blue line shows correlation between all points excluding the striatal FPCN, the most discriminatory feature (*r*=0.11, *p*=0.74). **(d)** ROC curves of a classifier trained with interaction effects between the two strongest-weighted seeds. In addition to the individual parcel features, we added the four-way interaction between the cerebellar FPCN and DAN and the striatal FPCN and DAN as features. Again, the classifier performance was not improved, indicating that no additional information is contributed by the interactive effect.

Collectively, we show that SVM binary classifiers using cerebellar and striatal connectivity features can distinguish between groups with high sensitivity and specificity. These classifier results are consistent with prior analyses indicating that neither cerebellum nor striatum as a whole drives classifier performance, suggesting that network disruptions in SCZ persists across both systems. Furthermore, associative parcels in both cerebellum and striatum are particularly discriminatory between groups, supporting both *a priori* and data-driven results presented above. That is, brain-wide disturbance in SCZ appear most pronounced for higher-order associative subdivisions across cerebellar and striatal functional parcels.

### Cerebellar and Striatal Dysconnectivity Relationship with Symptoms

While the observed effects reflected robust between-group differences, we also tested if CSTC alterations relate to symptom measures across subjects. Notably, there were no significant correlations between the degree of cerebellar or striatal dysconnectivity across patients (hyper-connectivity, hypo-connectivity, or a linear combination of the two) and SCZ symptom severity (as measured by the Positive and Negative Symptom Scale (PANSS)). This absence held for individual symptoms (**Supplementary Fig. S38-S39**) as well as when using the classic three-factor model of the Positive and Negative Syndrome Scale – a well-established measure of symptoms across the psychosis spectrum (PANSS, positive, negative, and general symptoms (Kay SR *et al.* 1987); see **Supplementary Fig. S40-S41**, *p*>0.05 with Bonferroni correction). To ensure a comprehensive characterization of symptom-related effects, we additionally investigated the relationship between identified dysconnectivity and an alternative five-factor model of PANSS, shown to be stable in an external sample of 5,789 subjects (van der Gaag M et al. 2006). These additional analyses again revealed weak correlations between the degree of dysconnectivity and the five-factor model symptom solution (**Supplementary Fig. S42-S43**). These comprehensive symptom analyses suggest that the identified CSTC dysconnectivity likely constitutes a trait-like effect, as opposed to a robust state-like marker, in line with prior observations that revealed modest relationships with thalamic connectivity (Anticevic A, MW Cole*, et al.* 2014). As such, these brain-wide differences do not reflect patients’ current symptom status, but may rather indicate a stable, disease-associated change.

### Cerebellar and Striatal Dysconnectivity Relationship with Cognition

Above we tested whether CSTC dysconnectivity relates to symptom measures. We observed weak or no statistical relationships despite adequate power. One alternative possibility is that the observed dysconnectivity may reflect cognitive deficits – a possibly in line with recent reports showing that thalamic connectivity may provide a cognitive remediation target in schizophrenia (Ramsay IS *et al.* 2017). To test this in relation to CSTC dysconnectivity, we first replicated the cerebellar and striatal connectivity analyses in an independent sample of 145 patients with schizophrenia and healthy controls from the Bipolar-Schizophrenia Network on Intermediate Phenotypes (B-SNIP) dataset, which also included measures of cognition (**Fig. 10**; see **Materials and Methods** and **Supplementary Tables S1-S2** for details) (Tamminga CA *et al.* 2013). We quantified the relationship between the degree of neural dysconnectivity and PANSS symptom severity as well as cognitive performance across individual subjects. As with the discovery sample, cerebellar and striatal dysconnectivity showed only weak correlations with symptom severity measures (**Supplementary Fig. S49-S54**). However, **Fig. 10c** shows a significant negative correlation between cerebellar hyper-connectivity and a measure of cognitive performance (the Brief Assessment of Cognition in Schizophrenia (BACS) summary score (Keefe RS *et al.* 2004)), indicating that individuals with a higher degree of hyper-connectivity performed more poorly on the BACS (*r*=-0.280, *p*=4.47e-8). Similarly, individuals with the greatest cerebellar hypo-connectivity also had the lowest BACS scores (*r*=0.301, *p*=3.62e-9, **Fig 10d**). These relationships held for striatal hyper-connectivity (*r*=-0.282, *p*=3.38e-8, **Fig. 10g**) and hypo-connectivity (*r*=0.271, *p*=1.17e-7, **Fig. 10h**). Notably, the preferentially stronger dysconnectivity with associative cerebellar and striatal subdivisions replicated (**Supplementary Fig. S44-S45**), and this FPCN dysconnectivity was significantly correlated with cognitive performance (**Supplementary Fig. S46** and **S48**). The relationship between cognition and CSTC dysconnectivity (either cerebellum or striatum for either hypo- or hyper-directions) accounted for close to 10% of variance across subjects, whereas the top relationships across all summary measures for symptom scores accounted for ∼2.3% of the variance (see **Supplementary Fig. S38-S43)**. This observation is consistent with the possibility that the brain-wide CSTC dysconnectivity reflects cognitive disturbances in SCZ.

**Figure 10.**
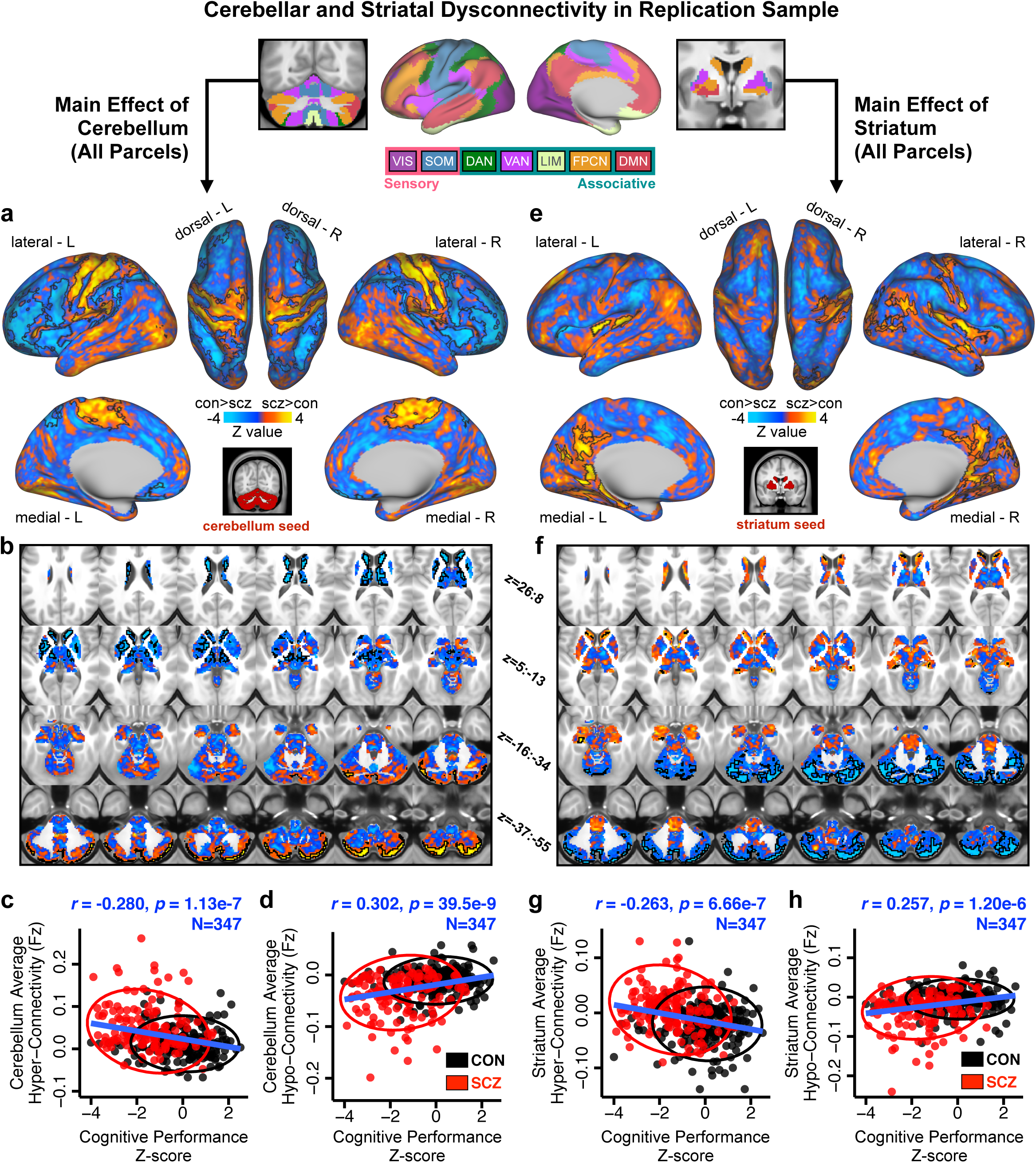
Main effect of cerebellar and striatal network parcel connectivity in replication sample and relationship with cognition. **(a)** Cortical surface view of areas showing main effect of *Group* in whole-brain connectivity with the cerebellar parcels between 145 patients with schizophrenia with psychosis (SCZ) and 202 healthy controls (CON) in an independent dataset. Regions that are significantly different between groups (*p*<0.05 with TFCE family-wise error protection) are outlined in black (see also **Supplementary Fig. S47**). Orange/yellow areas indicate regions where patients exhibited stronger cerebellar connectivity, whereas blue areas indicate regions where patients exhibited reduced cerebellar connectivity, relative to controls. Inset shows coverage of all cerebellar parcels. **(b)** Subcortical group differences in cerebellar connectivity shown in volume-based axial view, with Z-coordinate ranges (each slice in each row increments by 3 mm). **(c)** Relationship between mean cerebellar hyper-connectivity and a measure of cognitive performance is highly significant across subjects. (d) Relationship between mean cerebellar hypo-connectivity and cognitive performance is also highly significant across subjects. **(e-h)** Results for identical independent analysis conducted with the 6 functionally-defined striatal network parcels. Note that there was no representation of the visual network in striatum (Choi EY *et al.* 2012) and therefore that network was omitted. Abbreviations: VIS, visual network; SOM, somatosensory; DAN, dorsal attention network; VAN, ventral attention network; LIM, limbic network; FPCN, frontoparietal control network; DMN, default mode network.

## DISCUSSION

Identifying robust, replicable and neurobiologically-grounded neuroimaging markers for neuropsychiatric illness in general and SCZ in particular remains a major knowledge gap and an obstacle for rational treatment development. An ongoing debate concerns a tension between hypotheses suggesting that SCZ neural disturbances manifest across large-scale functional networks and those suggesting they occur in select regions (Anticevic A and J Lisman 2017). Informing this this question is important to facilitate biomarker development that can guide treatments for either targeted neuropathology in specific areas or diffuse alterations that span brain-wide pathways. We show that robust and replicable bi-directional dysconnectivity is present across CSTC, which were more pronounced in higher-order associative areas than in sensory areas. Using complementary data-driven approaches, we demonstrate that the cerebellum and striatum can be divided into functionally distinct disrupted circuits. Additionally, we demonstrate, using SVM-based machine learning, that ‘core’ associative alterations appear across both cerebellum and striatum and yield robust classification tools. Finally, we replicate the CSTC effects and we show that observed dysconnectivity weakly correlates with psychosis symptom severity but more strongly relates to cognitive traits across individual subjects. Collectively, these data are consistent with brain-wide bi-directional alterations in CSTC circuits in SCZ that manifest preferentially across associative neural networks.

### Brain-wide Disruptions Occur Across CSTC Functional Circuits in SCZ

Numerous lines of evidence implicate localized regional disruptions in SCZ. For instance, the dopamine hypothesis of SCZ strongly points to a striatal disruption in DA signaling. This is supported by the clinical potency of first-generation antipsychotics (and to some extent, second-generation “atypical” (Seeman P 2002)), whereby blockade of striatal dopamine D2 receptors correlates with therapeutic efficacy for psychotic symptoms (Seeman P et al. 1975; Seeman P and T Lee 1975). In humans, these receptors are primarily expressed in the dorsal striatum (Lee T and P Seeman 1980; Mackay AV et al. 1982; Howes OD et al. 2009). Complementary work centered on the DA hypothesis implicates hippocampal, thalamic and focal ventral tegmental area (VTA) alterations as central in the formation of positive symptoms of SCZ (Grace AA et al. 2007; Lodge DJ and AA Grace 2011). Conversely, glutamatergic hypotheses of SCZ, such as the prominent NMDAR hypo-function model, generally suggest a more widespread synaptic disruption centered on cortical microcircuits (Krystal JH et al. 2003; Gonzalez-Burgos G and DA Lewis 2012). Behaviorally, the SCZ spectrum is associated with broad deficits, including impaired executive cognitive functioning, deficits in perception, belief formation, as well as severe affective deficits marked by amotivation and anhedonia (Kay SR *et al.* 1987; Berenbaum H and TF Oltmanns 1992; Barch DM 2005; Stephan KE et al. 2009; Leitman DI et al. 2010; Calderone DJ et al. 2013). Collectively, it has been challenging to link such a broad-ranging behavioral disturbances across the SCZ spectrum to a single or a few punctate brain regions.

To date, neuroimaging studies have provided support for both ‘local hotspot’ and ‘widespread’ alterations, largely as a consequence of analytic method used. For instance, the thalamus has featured prominently in theoretical models of SCZ, both as a region of core putative disruption and part of a broader dysfunctional cortico-striato-thalamic loop, and as a locus of perturbation in functional coupling (Woodward ND *et al.* 2012; Anticevic A, MW Cole*, et al.* 2014; Anticevic A and J Lisman 2017). However, examining if such perturbations appear across neural systems, or if they are localized to select regions, necessitates the consideration of large-scale brain-wide network. We leveraged existing findings in healthy individuals that have identified large-scale functional networks across the cerebral cortex, basal ganglia, and cerebellum (Buckner RL *et al.* 2011; Yeo BT *et al.* 2011; Choi EY *et al.* 2012). If SCZ involves synaptic alterations that extend beyond thalamus then the bi-directional dysconnectivity seen with the thalamus may also be present across other systems as a shared pattern of disruption. Presented results combined complimentary analytic approaches, namely *a priori* subcortical network parcel seeds, voxelwise clustering of dysconnectivity, and binary classifiers of group status, to show that bi-directional disruptions in SCZ are indeed present across the striatum and cerebellum. Moreover, these alterations are particularly pronounced along associative higher-order functional subdivisions of these two systems. These three complementary results point to robust bi-directional alterations across both the striatum and cerebellum, extending more focused reports in SCZ subjects (Andreasen NC *et al.* 1996; Konarski JZ *et al.* 2005; Collin G *et al.* 2011; Tu PC *et al.* 2012; Fornito A et al. 2013; Sarpal DK *et al.* 2015; Horga G et al. 2016), as well as in related psychiatric disorders (Anand A et al. 2009; Harrison BJ et al. 2009; Cubillo A et al. 2010; Di Martino A et al. 2011; Anticevic A, S Hu, et al. 2014; Dandash O et al. 2014; Shinn AK et al. 2015).

### Treatment and Diagnostic Implications of Disturbances Across CSTC

While SCZ patients exhibited changes across both striatum and cerebellum, the associative higher-order subdivisions across the CSTC systems exhibited the most robust disturbances. That is, CSTC dysconnectivity in SCZ was most prominent along higher-order associative networks within both the cerebellum and striatum, in line with prior studies focused on thalamic and cortical networks (Barch DM et al. 2001; Camchong J et al. 2011; Woodward ND *et al.* 2012; Anticevic A, MW Cole*, et al.* 2014; Baker JT *et al.* 2014). This preferentially more severe associative disruption may reflect pervasive brain-wide synaptic deficits (e.g. NMDAR dysfunction), which in turn affects the most ‘vulnerable’ circuits (i.e. associative cortex). Recent biophysically-informed computational modeling work showed that widespread cellular-level glutamatergic deficits can yield preferentially more severe associative network dysconnectivity due to higher recurrent excitation in associative (versus sensory) cortex (Murray JD, A Bernacchia, et al. 2014; Scholtens LH et al. 2014; Yang GJ, JD Murray, XJ Wang*, et al.* 2016). Such differences in neuronal structure and functional coupling may extend throughout CSTC systems and may underlie the preferentially patterns of dysconnectivity observed across the cerebellum and striatum.

Critically, these findings present a challenge for treatment design. There appears to be some final converging regional expression of more severe dysconnectivity in association networks across cortex, thalamus, striatum and cerebellum. Consequently, do therapeutic effects need to occur *somewhere* (i.e. hotspot of disturbance if it indeed exists or emergent associative cortex disruption), *anywhere* (i.e. at any point of the affected CSTC circuit) or *everywhere* (i.e. global tuning of excitation/inhibition balance across all circuits), and at what stage of disease progression? A way to address this question is to longitudinally study duration-of-untreated psychosis (DUP) in early-course illness as a predictor of currently reported CSTC alterations and to map the neurobiology of at-risk clinical populations.

Notably, cerebellar and striatal features yielded high between-group single-subject classification accuracy (comparable to other classifiers in the literature (Davatzikos C et al. 2005; Kawasaki Y et al. 2007; Ardekani BA et al. 2011; Iwabuchi SJ et al. 2013; Schnack HG et al. 2014)). Nevertheless, there are key considerations for clinical applicability of the neuroimaging effects. First, the observed dysconnectivity effects did not relate to state-like measures of psychosis symptom severity, across several well-powered analyses for both the cerebellum and the striatum. This modest symptom effect was observed in the context of highly robust individual differences in hyper-/hypo-connectivity relationships across both the cerebellum and the striatum (see **Fig. 6**), which was replicated in an independent sample (i.e. not a floor or ceiling effect in neural data). Critically, in an independent similarly-sized SCZ replication sample we observed a significant relationship between cognition and CSTC dysconnectivity (either cerebellum or striatum for both hypo- or hyper-directions in cognition) across individuals. This is consistent with the possibility that CSTC dysconnectivity reflects cognitive disturbances as opposed to psychosis symptoms. These findings illustrate the complexity in generating the mapping between reliable group-level markers and clinically-meaningful behavioral dimensions. Future studies will need to develop a unified ‘geometry’ that integrates both categorical disturbances while concurrently capturing unique dimensions of clinical symptom variation.

Another key corollary of this idea is whether CSTC dysconnectivity appears across different psychiatric DSM-defined diagnoses, or if there are clear patterns that differentiate subgroups of individuals. Earlier results have shown that patients with bipolar disorder exhibit similar, but attenuated, bidirectional patterns of disruption in whole-brain thalamic dysconnectivity (Anticevic A, MW Cole*, et al.* 2014), with some unique features. Studying CSTC systems across diagnostic boundaries may reveal similarities in patterns of disruption, which may map onto shared disturbances across DSM-defined diagnoses (e.g. cognitive deficits).

Lastly, as noted, a key objective of this study was to test if cerebellar or striatal systems provide unique information for categorical single-subject classification. We used BOLD functional connectivity as the only dependent measure, showing that there are no major informational tradeoffs across the cerebellum or striatum. Instead, the classifiers performed best when aggregating information across associative networks across both systems. However, classifier accuracy could be improved by integrating information across multiple modalities that can map CSTC systems (Iwabuchi SJ *et al.* 2013), such as diffusion-weighted imaging (DWI). While each technique faces limitations, the combination of BOLD-DWI multimodal neuroimaging has the opportunity to be greater than the sum of its parts as it carries unique sources of information. Leveraging neuroimaging information across modalities may improve the accuracy and generalizability of classifiers across CSTC systems.

### Limitations, Pitfalls and Future Approaches

Importantly, while these results strongly indicate that disruptions in SCZ are pervasive across cerebellar and striatal systems, they do not rule out the possibility that these disturbances emerge over time as a consequence of a local ‘hotspot’ of disruption. Addressing this question will require prospective longitudinal clinical design that starts from the prodrome and examines if specific regional markers of dysfunction yield superior prospective classifier performance that in turn predict symptom severity. As discussed, leveraging diagnostically relevant information across different neuroimaging modalities may increase classifier performance in relation to state/trait predictive power. Another outstanding question concerns whether this observed dysconnectivity is consistent across different DSM-defined disorders. Studying dysconnectivity across CSTC systems using matched cross-diagnostic samples may reveal patterns of bi-directional CSTC disruption in some patients exclusively (i.e. DSM-like categorical pattern) or perhaps may appear in all patients as a function of cognitive deficit severity (RDoC-like dimensional pattern) (Insel T et al. 2010).

### Conclusion

SCZ research faces a tension between hypotheses proposing ‘focal hotspot’ disturbances affecting neural communication versus pervasive alterations that span distributed neural circuits. We demonstrate that similar bi-directional disruptions in functional coupling are robustly present across both cerebellar and striatal subdivisions of the CSTC system, with preferentially more pronounced alterations along ‘associative’ CSTC subdivisions. This shared and widespread CSTC dysconnectivity in SCZ is inconsistent with the possibility of exclusive ‘localized’ thalamic disruptions (at least in chronic states). Notably, results replicated across multiple complementary analytic approaches and independent datasets, linking alterations across CSTC systems with cognition in SCZ.

## ACKNOWLEDGEMENTS

Data were provided by the Centers of Biomedical Research Excellence (COBRE), the Olin Neuropsychiatric Research Center, and the Bipolar-Schizophrenia Network on Intermediate Phenotypes consortium.

## References

Anand A, Li Y, Wang Y, Lowe MJ, Dzemidzic M. 2009. Resting state corticolimbic connectivity abnormalities in unmedicated bipolar disorder and unipolar depression. Psychiatry Res 171:189–198.

Andreasen NC, O’Leary DS, Cizadlo T, Arndt S, Rezai K, Ponto LL, Watkins GL, Hichwa RD. 1996. Schizophrenia and cognitive dysmetria: a positron-emission tomography study of dysfunctional prefrontal-thalamic-cerebellar circuitry. Proc Natl Acad Sci U S A 93:9985–9990.

Andreasen NC, Pressler M, Nopoulos P, Miller D, Ho BC. 2010. Antipsychotic dose equivalents and dose-years: a standardized method for comparing exposure to different drugs. Biol Psychiatry 67:255–262.

Anticevic A, Brumbaugh MS, Winkler AM, Lombardo LE, Barrett J, Corlett PR, Kober H, Gruber J, Repovs G, Cole MW, Krystal JH, Pearlson GD, Glahn DC. 2012. Global Prefrontal and Fronto-amygdala Dysconnectivity in Bipolar I Disorder with Psychosis History. Biological Psychiatry 73:565–573.

Anticevic A, Cole MW, Repovs G, Murray JD, Brumbaugh MS, Winkler AM, Savic A, Krystal JH, Pearlson GD, Glahn DC. 2014. Characterizing thalamo-cortical disturbances in schizophrenia and bipolar illness. Cereb Cortex 24:3116–3130.

Anticevic A, Gancsos M, Murray JD, Repovs G, Driesen NR, Ennis DJ, Niciu MJ, Morgan PT, Surti TS, Bloch MH, Ramani R, Smith MA, Wang XJ, Krystal JH, Corlett PR. 2012. NMDA receptor function in large-scale anticorrelated neural systems with implications for cognition and schizophrenia. Proc Natl Acad Sci U S A 109:16720–16725.

Anticevic A, Haut K, Murray JD, Repovs G, Yang GJ, Diehl C, McEwen SC, Bearden CE, Addington J, Goodyear B, Cadenhead KS, Mirzakhanian H, Cornblatt BA, Olvet D, Mathalon DH, McGlashan TH, Perkins DO, Belger A, Seidman LJ, Tsuang MT, van Erp TG, Walker EF, Hamann S, Woods SW, Qiu M, Cannon TD. 2015. Association of Thalamic Dysconnectivity and Conversion to Psychosis in Youth and Young Adults at Elevated Clinical Risk. JAMA Psychiatry 72:882–891.

Anticevic A, Hu S, Zhang S, Savic A, Billingslea E, Wasylink S, Repovs G, Cole MW, Bednarski S, Krystal JH, Bloch MH, Li CS, Pittenger C. 2014. Global resting-state functional magnetic resonance imaging analysis identifies frontal cortex, striatal, and cerebellar dysconnectivity in obsessive-compulsive disorder. Biol Psychiatry 75:595–605.

Anticevic A, Lisman J. 2017. How Can Global Alteration of Excitation/Inhibition Balance Lead to the Local Dysfunctions That Underlie Schizophrenia? Biol Psychiatry.

Anticevic A, Repovs G, Barch DM. 2012. Emotion effects on attention, amygdala activation, and functional connectivity in schizophrenia. Schizophr Bull 38:967–980.

Ardekani BA, Tabesh A, Sevy S, Robinson DG, Bilder RM, Szeszko PR. 2011. Diffusion tensor imaging reliably differentiates patients with schizophrenia from healthy volunteers. Hum Brain Mapp 32:1–9.

Baker JT, Holmes AJ, Masters GA, Yeo BT, Krienen F, Buckner RL, Ongür D. 2014. Disruption of cortical association networks in schizophrenia and psychotic bipolar disorder. JAMA Psychiatry 72:109–118.

Barch DM. 2005. The relationships among cognition, motivation, and emotion in schizophrenia: how much and how little we know. Schizophrenia Bulletin 31:875–881.

Barch DM, Carter CS, Braver TS, Sabb FW, MacDonald A, 3rd, Noll DC, Cohen JD. 2001. Selective deficits in prefrontal cortex function in medication-naive patients with schizophrenia. Arch Gen Psychiatry 58:280–288.

Berenbaum H, Oltmanns TF. 1992. Emotional experience and expression in schizophrenia and depression. Journal of Abnormal Psychology 101:37–44.

Buckner RL, Krienen FM, Castellanos A, Diaz JC, Yeo BT. 2011. The organization of the human cerebellum estimated by intrinsic functional connectivity. J Neurophysiol 106:2322–2345.

Calderone DJ, Martinez A, Zemon V, Hoptman MJ, Hu G, Watkins JE, Javitt DC, Butler PD. 2013. Comparison of psychophysical, electrophysiological, and fMRI assessment of visual contrast responses in patients with schizophrenia. Neuroimage 67:153–162.

Camchong J, Macdonald AW, Bell C, Mueller BA, Lim KO. 2011. Altered functional and anatomical connectivity in schizophrenia. Schizophrenia bulletin 37:640–650.

Cauda F, D’Agata F, Sacco K, Duca S, Geminiani G, Vercelli A. 2011. Functional connectivity of the insula in the resting brain. Neuroimage 55:8–23.

Choi EY, Yeo BT, Buckner RL. 2012. The organization of the human striatum estimated by intrinsic functional connectivity. J Neurophysiol 108:2242–2263.

Clopper CJ, Pearson ES. 1934. The Use of Confidence or Fiducial Limits Illustrated in the Case of the Binomial. Biometrika 26:404–413.

Cole MW, Anticevic A, Repovs G, Barch DM. 2011. Variable global dysconnectivity and individual differences in schizophrenia. Biological Psychiatry 70:43–50.

Collin G, Hulshoff Pol HE, Haijma SV, Cahn W, Kahn RS, van den Heuvel MP. 2011. Impaired cerebellar functional connectivity in schizophrenia patients and their healthy siblings. Front Psychiatry 2:73.

Cubillo A, Halari R, Ecker C, Giampietro V, Taylor E, Rubia K. 2010. Reduced activation and inter-regional functional connectivity of fronto-striatal networks in adults with childhood Attention-Deficit Hyperactivity Disorder (ADHD) and persisting symptoms during tasks of motor inhibition and cognitive switching. J Psychiatr Res 44:629–639.

Dandash O, Fornito A, Lee J, Keefe RS, Chee MW, Adcock RA, Pantelis C, Wood SJ, Harrison BJ. 2014. Altered striatal functional connectivity in subjects with an at-risk mental state for psychosis. Schizophr Bull 40:904–913.

Davatzikos C, Shen D, Gur RC, Wu X, Liu D, Fan Y, Hughett P, Turetsky BI, Gur RE. 2005. Whole-brain morphometric study of schizophrenia revealing a spatially complex set of focal abnormalities. Arch Gen Psychiatry 62:1218–1227.

Di Martino A, Kelly C, Grzadzinski R, Zuo XN, Mennes M, Mairena MA, Lord C, Castellanos FX, Milham MP. 2011. Aberrant striatal functional connectivity in children with autism. Biol Psychiatry 69:847–856.

Fornito A, Harrison BJ, Goodby E, Dean A, Ooi C, Nathan PJ, Lennox BR, Jones PB, Suckling J, Bullmore ET. 2013. Functional Dysconnectivity of Corticostriatal Circuitry as a Risk Phenotype for Psychosis. JAMA psychiatry (Chicago, Ill).

Glasser MF, Sotiropoulos SN, Wilson JA, Coalson TS, Fischl B, Andersson JL, Xu J, Jbabdi S, Webster M, Polimeni JR, Van Essen DC, Jenkinson M, Consortium WU-MH. 2013. The minimal preprocessing pipelines for the Human Connectome Project. Neuroimage 80:105–124.

Gonzalez-Burgos G, Lewis DA. 2012. NMDA Receptor Hypofunction, Parvalbumin-Positive Neurons and Cortical Gamma Oscillations in Schizophrenia. Schizophrenia Bulletin 38:950–957.

Grace AA, Floresco SB, Goto Y, Lodge DJ. 2007. Regulation of firing of dopaminergic neurons and control of goal-directed behaviors. Trends Neurosci 30:220–227.

Haber S, McFarland NR. 2001. The place of the thalamus in frontal cortical-basal ganglia circuits. Neuroscientist 7:315–324.

Harrison BJ, Soriano-Mas C, Pujol J, Ortiz H, López-Solà M, Hernández-Ribas R, Deus J, Alonso P, Yücel M, Pantelis C, Menchon JM, Cardoner N. 2009. Altered corticostriatal functional connectivity in obsessive-compulsive disorder. Archives of General Psychiatry 66:1189–1200.

Holt DJ, Bachus SE, Hyde TM, Wittie M, Herman MM, Vangel M, Saper CB, Kleinman JE. 2005. Reduced density of cholinergic interneurons in the ventral striatum in schizophrenia: an in situ hybridization study. Biol Psychiatry 58:408–416.

Horga G, Cassidy CM, Xu X, Moore H, Slifstein M, Van Snellenberg JX, Abi-Dargham A. 2016. Dopamine-Related Disruption of Functional Topography of Striatal Connections in Unmedicated Patients With Schizophrenia. JAMA Psychiatry 73:862–870.

Howes OD, Montgomery AJ, Asselin MC, Murray RM, Valli I, Tabraham P, Bramon-Bosch E, Valmaggia L, Johns L, Broome M, McGuire PK, Grasby PM. 2009. Elevated striatal dopamine function linked to prodromal signs of schizophrenia. Arch Gen Psychiatry 66:13–20.

Insel T, Cuthbert B, Garvey M, Heinssen R, Pine DS, Quinn K, Sanislow C, Wang P. 2010. Research domain criteria (RDoC): toward a new classification framework for research on mental disorders. Am J Psychiatry 167:748–751.

Iwabuchi SJ, Liddle PF, Palaniyappan L. 2013. Clinical utility of machine-learning approaches in schizophrenia: improving diagnostic confidence for translational neuroimaging. Frontiers in psychiatry 4:95.

Kawasaki Y, Suzuki M, Kherif F, Takahashi T, Zhou SY, Nakamura K, Matsui M, Sumiyoshi T, Seto H, Kurachi M. 2007. Multivariate voxel-based morphometry successfully differentiates schizophrenia patients from healthy controls. Neuroimage 34:235–242.

Kay SR, Fiszbein A, Opler LA. 1987. The positive and negative syndrome scale (PANSS) for schizophrenia. Schizophrenia Bulletin 13:261–276.

Keefe RS, Goldberg TE, Harvey PD, Gold JM, Poe MP, Coughenour L. 2004. The Brief Assessment of Cognition in Schizophrenia: reliability, sensitivity, and comparison with a standard neurocognitive battery. Schizophr Res 68:283–297.

Klein JC, Rushworth MFS, Behrens TEJ, Mackay CE, de Crespigny AJ, D’Arceuil H, Johansen-Berg H. 2010. Topography of connections between human prefrontal cortex and mediodorsal thalamus studied with diffusion tractography. NeuroImage 51:555–564.

Konarski JZ, McIntyre RS, Grupp LA, Kennedy SH. 2005. Is the cerebellum relevant in the circuitry of neuropsychiatric disorders? J Psychiatry Neurosci 30:178–186.

Krystal JH, D’Souza DC, Mathalon D, Perry E, Belger A, Hoffman R. 2003. NMDA receptor antagonist effects, cortical glutamatergic function, and schizophrenia: toward a paradigm shift in medication development. Psychopharmacology 169:215–233.

Lee T, Seeman P. 1980. Elevation of brain neuroleptic/dopamine receptors in schizophrenia. Am J Psychiatry 137:191–197.

Leitman DI, Sehatpour P, Higgins BA, Foxe JJ, Silipo G, Javitt DC. 2010. Sensory deficits and distributed hierarchical dysfunction in schizophrenia. Am J Psychiatry 167:818–827.

Lisman JE, Pi HJ, Zhang Y, Otmakhova NA. 2010. A thalamo-hippocampal-ventral tegmental area loop may produce the positive feedback that underlies the psychotic break in schizophrenia. Biological Psychiatry 68:17–24.

Liu H, Fan G, Xu K, Wang F. 2011. Changes in cerebellar functional connectivity and anatomical connectivity in schizophrenia: a combined resting-state functional MRI and diffusion tensor imaging study. J Magn Reson Imaging 34:1430–1438.

Lodge DJ, Grace AA. 2011. Hippocampal dysregulation of dopamine system function and the pathophysiology of schizophrenia. Trends Pharmacol Sci 32:507–513.

Mackay AV, Iversen LL, Rossor M, Spokes E, Bird E, Arregui A, Creese I, Synder SH. 1982. Increased brain dopamine and dopamine receptors in schizophrenia. Arch Gen Psychiatry 39:991–997.

Meda SA, Wang Z, Ivleva EI, Poudyal G, Keshavan MS, Tamminga CA, Sweeney JA, Clementz BA, Schretlen DJ, Calhoun VD, Lui S, Damaraju E, Pearlson GD. 2015. Frequency-Specific Neural Signatures of Spontaneous Low-Frequency Resting State Fluctuations in Psychosis: Evidence From Bipolar-Schizophrenia Network on Intermediate Phenotypes (B-SNIP) Consortium. Schizophr Bull 41:1336–1348.

Meyer-Lindenberg AS, Olsen RK, Kohn PD, Brown T, Egan MF, Weinberger DR, Berman KF. 2005. Regionally specific disturbance of dorsolateral prefrontal-hippocampal functional connectivity in schizophrenia. Arch Gen Psychiatry 62:379–386.

Murray JD, Anticevic A, Gancsos M, Ichinose M, Corlett PR, Krystal JH, Wang XJ. 2014. Linking microcircuit dysfunction to cognitive impairment: effects of disinhibition associated with schizophrenia in a cortical working memory model. Cereb Cortex 24:859–872.

Murray JD, Bernacchia A, Freedman DJ, Romo R, Wallis JD, Cai X, Padoa-Schioppa C, Pasternak T, Seo H, Lee D, Wang XJ. 2014. A hierarchy of intrinsic timescales across primate cortex. Nat Neurosci 17:1661–1663.

Nanetti L, Cerliani L, Gazzola V, Renken R, Keysers C. 2009. Group analyses of connectivity-based cortical parcellation using repeated k-means clustering. Neuroimage 47:1666–1677.

Power JD, Barnes KA, Snyder AZ, Schlaggar BL, Petersen SE. 2012. Steps toward optimizing motion artifact removal in functional connectivity MRI; a reply to Carp. Neuroimage 76:439–441.

Ramsay IS, MacDonald AW, 3rd. 2018. The Ups and Downs of Thalamocortical Connectivity in Schizophrenia. Biol Psychiatry 83:473–474.

Ramsay IS, Nienow TM, MacDonald AW, 3rd. 2017. Increases in Intrinsic Thalamocortical Connectivity and Overall Cognition Following Cognitive Remediation in Chronic Schizophrenia. Biol Psychiatry Cogn Neurosci Neuroimaging 2:355–362.

Ray JP, Price JL. 1993. The organization of projections from the mediodorsal nucleus of the thalamus to orbital and medial prefrontal cortex in macaque monkeys. Journal of Comparative Neurology 337:1–31.

Repovs G, Barch DM. 2012. Working memory related brain network connectivity in individuals with schizophrenia and their siblings. Frontiers in Human Neuroscience 6.

Repovs G, Csernansky JG, Barch DM. 2011. Brain network connectivity in individuals with schizophrenia and their siblings. Biological Psychiatry 15:967–973.

Rusch N, Spoletini I, Wilke M, Bria P, Di Paola M, Di Iulio F, Martinotti G, Caltagirone C, Spalletta G. 2007. Prefrontal-thalamic-cerebellar gray matter networks and executive functioning in schizophrenia. Schizophr Res 93:79–89.

Sarpal DK, Argyelan M, Robinson DG, Szeszko PR, Karlsgodt KH, John M, Weissman N, Gallego JA, Kane JM, Lencz T, Malhotra AK. 2016. Baseline Striatal Functional Connectivity as a Predictor of Response to Antipsychotic Drug Treatment. American Journal of Psychiatry 173:69–77.

Sarpal DK, Robinson DG, Lencz T, Argyelan M, Ikuta T, Karlsgodt KH, Gallego JA, Kane JM, Szeszko PR, Malhotra AK. 2015. Antipsychotic treatment and functional connectivity of the striatum in first-episode schizophrenia. JAMA Psychiatry 72:5–13.

Schnack HG, Nieuwenhuis M, van Haren NE, Abramovic L, Scheewe TW, Brouwer RM, Hulshoff Pol HE, Kahn RS. 2014. Can structural MRI aid in clinical classification? A machine learning study in two independent samples of patients with schizophrenia, bipolar disorder and healthy subjects. Neuroimage 84:299–306.

Scholtens LH, Schmidt R, de Reus MA, van den Heuvel MP. 2014. Linking macroscale graph analytical organization to microscale neuroarchitectonics in the macaque connectome. J Neurosci 34:12192–12205.

Seeman P. 2002. Atypical antipsychotics: mechanism of action. Can J Psychiatry 47:27–38.

Seeman P, Chau-Wong M, Tedesco J, Wong K. 1975. Brain receptors for antipsychotic drugs and dopamine: direct binding assays. Proc Natl Acad Sci U S A 72:4376–4380.

Seeman P, Lee T. 1975. Antipsychotic drugs: direct correlation between clinical potency and presynaptic action on dopamine neurons. Science 188:1217–1219.

Sheffield JM, Kandala S, Tamminga CA, Pearlson GD, Keshavan MS, Sweeney JA, Clementz BA, Lerman-Sinkoff DB, Hill SK, Barch DM. 2017. Transdiagnostic Associations Between Functional Brain Network Integrity and Cognition. JAMA Psychiatry 74:605–613.

Shinn AK, Baker JT, Lewandowski KE, Ongur D, Cohen BM. 2015. Aberrant cerebellar connectivity in motor and association networks in schizophrenia. Front Hum Neurosci 9:134.

Smith SM, Nichols TE. 2009. Threshold-free cluster enhancement: addressing problems of smoothing, threshold dependence and localisation in cluster inference. Neuroimage 44:83–98.

Stephan KE, Baldeweg T, Friston KJ. 2006. Synaptic plasticity and dysconnection in schizophrenia. Biol Psychiatry 59:929–939.

Stephan KE, Friston KJ, Frith CD. 2009. Dysconnection in schizophrenia: from abnormal synaptic plasticity to failures of self-monitoring. Schizophrenia bulletin 35:509–527.

Tamminga CA, Ivleva EI, Keshavan MS, Pearlson GD, Clementz BA, Witte B, Morris DW, Bishop J, Thaker GK, Sweeney JA. 2013. Clinical phenotypes of psychosis in the Bipolar-Schizophrenia Network on Intermediate Phenotypes (B-SNIP). Am J Psychiatry 170:1263–1274.

Tu PC, Hsieh JC, Li CT, Bai YM, Su TP. 2012. Cortico-striatal disconnection within the cingulo-opercular network in schizophrenia revealed by intrinsic functional connectivity analysis: A resting fMRI study. NeuroImage.

van der Gaag M, Hoffman T, Remijsen M, Hijman R, de Haan L, van Meijel B, van Harten PN, Valmaggia L, de Hert M, Cuijpers A, Wiersma D. 2006. The five-factor model of the Positive and Negative Syndrome Scale II: a ten-fold cross-validation of a revised model. Schizophr Res 85:280–287.

Wang L, Zou F, Shao Y, Ye E, Jin X, Tan S, Hu D, Yang Z. 2014. Disruptive changes of cerebellar functional connectivity with the default mode network in schizophrenia. Schizophr Res 160:67–72.

Welsh RC, Chen AC, Taylor SF. 2010. Low-frequency BOLD fluctuations demonstrate altered thalamocortical connectivity in schizophrenia. Schizophr Bull 36:713–722.

Winkler AM, Ridgway GR, Douaud G, Nichols TE, Smith SM. 2016. Faster permutation inference in brain imaging. Neuroimage 141:502–516.

Winkler AM, Ridgway GR, Webster MA, Smith SM, Nichols TE. 2014. Permutation inference for the general linear model. Neuroimage 92:381–397.

Woodward ND, Karbasforoushan H, Heckers S. 2012. Thalamocortical dysconnectivity in schizophrenia. Am J Psychiatry 169:1092–1099.

Yang GJ, Murray JD, Wang X-J, Glahn DC, Pearlson GD, Repovs G, Krystal JH, Anticevic A. 2016. Functional hierarchy underlies preferential connectivity disturbances in schizophrenia. Proceedings of the National Academy of Science USA 113:E219–228.

Yang GJ, Murray JD, Wang XJ, Glahn DC, Pearlson GD, Repovs G, Krystal JH, Anticevic A. 2016. Functional hierarchy underlies preferential connectivity disturbances in schizophrenia. Proc Natl Acad Sci U S A 113:E219–228.

Yeo BT, Krienen FM, Sepulcre J, Sabuncu MR, Lashkari D, Hollinshead M, Roffman JL, Smoller JW, Zollei L, Polimeni JR, Fischl B, Liu H, Buckner RL. 2011. The organization of the human cerebral cortex estimated by intrinsic functional connectivity. J Neurophysiol 106:1125–1165.

